# Genetic basis of growth, phenology and susceptibility to biotic stressors in maritime pine

**DOI:** 10.1101/699389

**Authors:** Agathe Hurel, Marina de Miguel, Cyril Dutech, Marie-Laure Desprez-Loustau, Christophe Plomion, Isabel Rodríguez-Quilón, Agathe Cyrille, Thomas Guzman, Ricardo Alía, Santiago C. González-Martínez, Katharina B. Budde

**Affiliations:** BIOGECO, INRAE, Univ. Bordeaux, 33610 Cestas, France; EGFV, INRAE, Univ. Bordeaux, 33140 Villenave-d’Ornon, France; CIFOR, INIA, 28040 Madrid, Spain; Georg-August University Göttingen, Büsgen-Institute, 37077 Göttingen, Germany

**Keywords:** *Pinus pinaster*, growth phenology, pathogen susceptibility, heritability, genetic correlations, association genetics

## Abstract

Forest ecosystems are increasingly challenged by extreme events, e.g. drought, storms, pest and pathogenic fungi outbreaks, causing severe ecological and economical losses. Understanding the genetic basis of adaptive traits in tree species is of key importance to preserve forest ecosystems, as genetic variation in a trait (i.e. heritability) determines its potential for human-mediated or evolutionary change. Maritime pine (*Pinus pinaster* Aiton), a conifer widely distributed in southwestern Europe and northwestern Africa, grows under contrasted environmental conditions promoting local adaptation. Genetic variation at adaptive phenotypes, including height, growth phenology and susceptibility to two fungal pathogens (*Diplodia sapinea* and *Armillaria ostoyae*) and an insect pest (*Thaumetopoea pityocampa*), were assessed in a range-wide clonal common garden of maritime pine. Broad-sense heritability was significant for height (0.219), growth phenology (0.165-0.310) and pathogen susceptibility (necrosis length caused by *D. sapinea*, 0.152; and by *A. ostoyae*, 0.021) measured after inoculation under controlled conditions, but not for pine processionary moth incidence in the common garden. The correlations of trait variation among populations revealed contrasting trends for pathogen susceptibility to *D. sapinea* and *A. ostoyae* with respect to height. Taller trees showed longer necrosis length caused by *D. sapinea* while shorter trees were more affected by *A. ostoyae*. Moreover, maritime pine populations from areas with high summer temperatures and frequent droughts were less susceptible to *D. sapinea* but more susceptible to *A. ostoyae*. Finally, an association study using 4,227 genome-wide SNPs revealed several loci significantly associated to each trait (range of 3-26), including a possibly disease-induced translation initiation factor, eIF-5. This study provides important insights to develop genetic conservation and breeding strategies integrating species responses to biotic stressors.

## Introduction

Forest ecosystems are challenged worldwide by changing environmental conditions (Turner, 2010). Warmer and drier climates are expected to increase fire risk, droughts and insect outbreaks while warmer and wetter climates will probably increase storm and pathogen incidence on forests (Seidl *et al*., 2017), leading to episodes of high tree mortality (Allen *et al*., 2010) and severe economic losses (Hanewinkel *et al*., 2013). Changing environmental conditions can also cause range shifts in previously locally restricted pests and pathogens or shifts to increased pathogenicity (Desprez-Loustau *et al*., 2006). Thus, understanding forest tree genetic variation in disease (and other stressor) responses, in relationship to growth and phenology, is crucial to develop informed forest restoration, conservation and management strategies. Moreover, genes underlying adaptive traits, such as growth or disease response, can serve tree breeding and increase forest productivity, e.g. targeting resistance to drought or against pathogens in forest plantations (Neale & Kremer, 2011).

Forest trees are long-lived, sessile organisms. They are characterized by outcrossing mating systems, high standing genetic variation, large effective population sizes, and the production of vast numbers of seeds and seedlings exposed to strong selection (Petit *et al*., 2004; Petit & Hampe, 2006). High genetic and phenotypic differentiation has been observed in tree species along environmental gradients (e.g. Savolainen *et al*., 2007, 2013) or between contrasting habitats, indicating local adaptation (e.g. Lind *et al*., 2017). Common garden experiments (i.e. experiments evaluating trees from a wide range of populations under the same environmental conditions) provide valuable insights on the phenotypic and genotypic variation of forest trees (Morgenstern, 2011). They revealed genetic differentiation for adaptive traits (such as flushing, senescence or growth) along latitudinal and altitudinal gradients (Mimura & Aitken, 2007; Vitasse *et al*., 2009). Geographical variation has also been found for disease resistance against certain pests (Menéndez-Gutiérrez *et al*., 2017) and pathogens (e.g. Hamilton *et al*., 2013; Freeman *et al*., 2019). Phenological traits, such as time to flowering, or spring bud burst and autumn leaf senescence times, are often genetically correlated with disease susceptibility, providing hints on the different kind of resistance or avoidance mechanisms found in forest trees (Elzinga *et al*., 2007).

Disease resistance is generally thought to be the result of selective pressures exerted by the pathogen, in areas where host and pathogen have co-existed during considerable periods of time, under the co-evolution hypothesis (e.g. Burdon & Thrall, 2000; Ennos, 2015). In this line, geographical variation in disease resistance has been interpreted in some cases as a result of past heterogeneous pathogen pressures within the range of a given host species (Ennos, 2015; Perry *et al*., 2016). However, the past distribution of pathogen species is often unknown (Desprez-Loustau *et al*., 2016), therefore, other processes than co-evolution, such as “ecological fitting” or “exaptation” should not be excluded (Agosta & Klemens, 2008). These biological processes have been suggested when, for example, variability in disease resistance was observed in tree species with no co-evolutionary history with a pathogen (Leimu & Koricheva, 2006; Freeman *et al*., 2019). Such resistance may have evolved in response to other pathogens but had broad-range efficacy, even to a novel pathogen. Generic mechanisms of resistance in conifers include the production of large amounts of non-volatile compounds (resin acids) that can act as mechanical barriers to infections (Shain, 1967; Phillips & Croteau, 1999) and volatile compounds (such as monoterpenes or phenols) that can be toxic to fungi (Cobb *et al*., 1968; Chou & Zabkiewicz, 1976). The composition of secondary metabolites can show marked geographic variation in tree species (Meijón *et al*., 2016). The evolution of plant defences against biotic stressors can also be shaped by differences in resource availability and environmental constraints throughout the host’s species distribution. Depending on resource availability, plants have evolved distinct strategies by investing either more in growth, to increase competition ability, or more in chemical and structural defences, to better respond to herbivores and pathogens (Herms & Mattson, 2004). Typically, faster growing trees invest more in inducible defences while slow growing trees invest more in constitutive defences (Moreira *et al*., 2014).

Many quantitative traits in forest species, including height, growth phenology and disease resistance, show significant heritability and often stronger differentiation (*Q*_ST_) among populations than neutral genetic markers (*F*_ST_) (Hamilton *et al*., 2013; see review in Lind *et al*., 2018), a common indication of adaptive divergence. Major-effect genes for growth, e.g. *korrigan* in *Pinus pinaster* (Cabezas *et al*., 2015), and resistance genes against forest pathogens, e.g. against the fusiform rust disease in *Pinus taeda* (Kuhlman *et al*., 2002) and against white pine blister rust in several other North American pine species (Sniezko, 2008; Weiss *et al*., 2020), have been identified in forest trees. However, most adaptive traits have a highly polygenic basis of quantitative inheritance, typically involving many loci with rather small effects (Goldfarb *et al*., 2013; De la Torre *et al*., 2019). Most association genetic studies in forest trees focused on wood property and growth traits to assist tree breeding (e.g. Pot *et al*., 2005; Neale *et al*., 2006; Beaulieu *et al*., 2011). Also, genetic association approaches identified some loci associated to other ecologically important traits, such as cold hardiness (e.g. Eckert *et al*., 2009; Holliday *et al*., 2010), drought tolerance (reviewed in Moran *et al*., 2017), or disease resistance (e.g. Liu *et al*., 2014; Resende *et al*., 2017). Compared to more intensively studied traits, association studies addressing biotic interaction traits, including responses to pests and pathogenic fungi, are still limited to few tree species.

Our study focused on maritime pine (*Pinus pinaster* Aiton, Pinaceae), a long-lived conifer from southwestern Europe and northern Africa with a wide ecological amplitude. We assessed height, growth phenology (bud burst and duration of bud burst) and susceptibility to pests/pathogens in a clonal common garden, which allowed us to explore variation in disease response and genetic correlations with other traits in range-wide populations of maritime pine. Considering disease and growth traits together is relevant from an evolutionary and ecological perspective, and can have important implications in terms of management, especially in breeding programs. We selected three important disease agents: two fungal pathogens, *Diplodia sapinea* (Botryosphaeriaceae) and *Armillaria ostoyae* (Physalacriaceae), as well as the pine processionary moth, *Thaumetopoea pityocampa* (Thaumetopoeidae), a main defoliator of pine forests.

Maritime pine has a highly fragmented natural range in the western Mediterranean Basin, the Atlas Mountains in Morocco, the Atlantic coast of southern France and the west coast of the Iberian Peninsula, and grows from sea level to over 2,000 m altitude. Genetic diversity and population structure is high in natural populations of maritime pine, especially in the Iberian Peninsula, possibly due to its long-term persistence in this region (Petit *et al*., 1995; Salvador *et al*., 2000; Bucci *et al*., 2007; Jaramillo-Correa *et al*., 2015). In addition, traits, such as stem form, height (González-Martínez *et al*., 2002), metabolite content (Meijón *et al*., 2016), drought response (Aranda *et al*., 2010; Gaspar *et al*., 2013) and pest and disease resistance (Schvester, 1982; Desprez-Loustau & Baradat, 1991; Burban *et al*., 1999; Elvira-Recuenco *et al*., 2014), are highly variable in maritime pine and often strongly differentiated across geographical regions. Maritime pine has also been widely planted and is currently exploited for timber and paper, for example, covering ~1.03 million ha in in southwestern France (including the Landes region), one of the largest plantation forests in Europe (Memento FCBA 2020, https://www.fcba.fr/ressources/memento-2020/). Despite the ecological and economical importance of maritime pine natural forests and plantations, only a few genetic association studies have been developed in this species. Lepoittevin *et al*. (2012) identified two loci associated to growth and wood cellulose content, respectively. Cabezas *et al*. (2015) revealed four SNPs in *korrigan* (a pine gene orthologous to an Arabidopsis degrading enzyme cellulase) also significantly associated to growth traits (total height and polycyclism). Finally, Bartholomé *et al*. (2016) reported four loci for stem straightness and three loci for height growth. In addition, Budde *et al*. (2014) were able to predict 29% of the phenotypic variation in a fire adaptive trait (proportion of serotinous cones) in eastern Spain based on 17 SNP loci. However, none of these studies targeted phenology or biotic interaction traits, such as disease resistance.

*Diplodia sapinea* is the causal agent of several diseases, such as tip-blight, canker or root collar necrosis in needles, shoots, stems and roots of conifers, eventually leading to mortality in case of severe attacks (Piou *et al*., 1991; Luchi *et al*., 2014). The pathogenicity of *D. sapinea* is associated to environmental conditions. It can remain in an endophytic form, i.e. without causing any symptoms, until stressful environmental conditions, such as drought (Stanosz *et al*., 2002; Desprez-Loustau *et al*., 2006), hail storms (Zwolinski *et al*., 1990), or changes in the nitrogen concentration of the soil (Piou *et al*., 1991; Stanosz *et al*., 2004), weaken the host and trigger *D. sapinea* pathogenicity. Trees from all ages are affected (Chou, 1978; Georgieva & Hlebarska, 2017), though seedlings and old trees show increased susceptibility (Swart & Wingfield, 1991). The fungus can be found in many conifers, especially in the genus *Pinus*. Iturritxa *et al*. (2013) classified maritime pine as moderately susceptible. The species was first described in Europe in 1823 under the name *Sphaeria sapinea*, and then received many synonyms (Piou *et al*., 1991). Recent surveys showed that *D. sapinea* is currently broadly distributed in all pine forests throughout the world, though its origin is debated (Burgess *et al*., 2004; Brodde *et al*., 2019). Serious damage associated with *D. sapinea* in Europe has only been reported in the last decades but it may become a serious threat to pine forests, as climate change will certainly favor pathogen activity by increasing temperature, and the frequency and intensity of drought events (Woolhouse *et al*., 2005; Desprez-Loustau *et al*., 2006; Boutte, 2018). In this line, recent outbreaks associated with *D. sapinea* in northern Europe suggest an ongoing northward expansion (Brodde *et al*., 2019).

*Armillaria ostoyae* is a root pathogen that causes white rot and butt rot disease in conifers, leading to growth deprivation, high mortality and major losses in timber wood; hence, its economic importance (Cruickshank, 2011; Heinzelmann *et al*., 2019). The species can be traced back to six millions years ago, both in Eurasia and North America (Tsykun *et al*., 2013; Koch *et al*., 2017). *Armillaria ostoyae* has been reported in all the conifer forests of the Northern Hemisphere but it appears to be replaced by *A. mellea* (Marxmüller & Guillaumin, 2005) in the Mediterranean due to higher temperatures and drought. It is likely that *A. ostoyae* would have co-existed for a long time with maritime pine in Europe (Tsykun *et al*., 2013) and consequently it could have been affected by the same extinction-recolonization events associated to past climatic changes as its host (Labbé *et al*., 2017a). *Armillaria ostoyae* is one of the most common fungal species in maritime pine forests, being particularly dangerous as it can act as both a parasite and a saprophyte (Cruickshank *et al*., 1997; Labbé *et al*., 2017b), i.e. the death of its host does not prevent its spread. In maritime pine, the severity of *A. ostoyae* symptoms is related to host age, with higher mortality in young trees (Lung-Escarmant & Guyon, 2004; Labbé *et al*., 2015). Climate change is predicted to foster an increased impact of *A. ostoyae* on conifer forests in the coming years (Kubiak *et al*., 2017).

*Thaumetopoea pityocampa* is considered the most severe defoliator insect in pine forests in southern Europe and northern Africa (Jactel *et al*., 2015), leading to severe growth loss (Jacquet *et al*., 2013). The species typically reproduces in summer followed by larval development during autumn and winter. Caterpillars and moths of *T. pityocampa* are sensitive to climatic and environmental conditions, and this pest is expected to expand its range following events of climate warming (Battisti *et al*., 2006; Toïgo *et al*., 2017).

The specific objectives of our study are to: 1) estimate phenotypic variability and heritability within and among range-wide populations of maritime pine for height, growth phenology and susceptibility to pests/pathogens; 2) test for adaptive divergence across the maritime pine range for these traits (i.e. *Q*_ST_ vs. *F*_ST_ approach); 3) analyze the pattern of trait correlation within and among populations, in particular to identify putative correlations that could be useful/detrimental for conservation and breeding programs; and 4) identify loci associated to disease-related, growth and phenology traits using genotype-phenotype association. Altogether, our approach, combining the evaluation of a clonal common garden and a genotyping array, produced relevant insights on the evolution, genetic basis, and architecture of adaptive traits in maritime pine, an ecologically and economically important forest tree species.

## Material and Methods

### Plant material and common garden measurements

A clonal common garden (CLONAPIN) was planted in 2011 in Cestas, southwestern France (for details see Rodríguez-Quilón, 2017). It includes trees from 35 populations of maritime pine covering the whole natural species distribution (see Table S1.1, Supporting Information for number of individuals and genotypes, and population coordinates of 32 populations included in this study), representing all known differentiated gene pools in the species (Central Spain, Southeastern Spain, Iberian Atlantic, French Atlantic, Corsica and Morocco; see Jaramillo-Correa *et al*., 2015). The common garden design consisted of eight randomized complete blocks, with one clonal copy (ramet) of each genotype replicated in each block. For the pathogen inoculation experiments, we chose genotypes from populations representing each of the six gene pools. Because of the higher logistical effort involved in inoculation protocols with respect to other measurements (see below), it was not possible to include all common garden genotypes in these experiments (see Table S1.1, Supporting Information).

Height, growth phenology (bud burst and duration of bud burst), and incidence of processionary moth (*T. pityocampa*), were measured in all individuals from 5-8 blocks, depending on the trait (sample size of 1,392-3,204 trees, see Table S1.1, Supporting Information). Pathogen susceptibility was assessed in a subset of genotypes using excised branches collected from the clonal trial (sample size of 180-453 branches, see Table S1.1, Supporting Information and below). Tree height was measured in 2015, four years after the establishment of the trial. Bud burst stage was evaluated using a phenological scale ranging from 0 to 5 (0: bud without elongation during winter, 1: elongation of the bud, 2: emergence of brachyblast, 3: brachyblast begins to space, 4: elongation of the needles, 5: total elongation of the needles; see Figure S2.1, Supporting Information). The Julian day of entry in each stage (S1 to S5) was scored for each tree. Julian days were converted into accumulated degree-days (0°C basis) from the first day of the year, to take into account the between year variability in temperature. The number of degree-days between stages 1 and 4 defines the duration of bud burst. Both growth phenology phenotypes, bud burst and duration of bud burst, were assessed in 2015 and 2017. The presence or absence of pine processionary moth nests (*T. pityocampa*) in the tree crowns was assessed in March 2018.

### Experimental evaluation of susceptibility to Diplodia sapinea

Inoculations were carried out on excised shoots taken from pines in the common garden (for a detailed laboratory protocol see Supporting Information S3.1). We used the pathogen strain Pier4, isolated from *P. nigra* cones in Pierroton, France (May 2017), and maintained on malt-agar medium. The identity of this strain as *D. sapinea*, was confirmed by sequencing the ITS region, amplified using the primers ITS1-F and ITS4 (Gardes & Bruns, 1993), and blasting it against the NCBI nucleotide database (Benoît Laurent, personal communication). Only current-year shoots at the phenological stages 3 to 5, i.e. with fully elongated buds but not fully mature, were collected, otherwise sampled randomly (see Supporting Information S2.1). For the inoculation, we removed a needle fascicle in the middle of each shoot with a scalpel. A 5 mm diameter plug of malt-agar taken at the active margin of a *D. sapinea* culture was put on the wound, mycelium-side down, and then wrapped in cellophane. Control shoots were treated in the same manner but with plugs of sterile rather than colonized malt-agar. The shoots were put in water and kept in a climatic chamber set at 20°C with a daily cycle of 12h of light and 12h of dark (Blodgett & Bonello, 2003; Iturritxa *et al*., 2013). Six days after the inoculation, we removed the cellophane and measured the lesion length around the inoculation point with a caliper. The shoots were not lignified and the lesions were visible. However, the surface was superficially stripped to see the limit of the lesion when it was not visible otherwise. Needle discoloration was also observed, and evaluated using a scale from 0: no discoloration to 3: all needles along the necrosis showed discoloration (see Figure S3.1, Supporting Information). To confirm that discoloration was caused by the pathogen, one discolored needle from one branch per population was placed on a malt-agar Petri dish to grow. After 3 days, *D. sapinea* could be visually identified in each Petri dish.

For this experiment, we sampled a total of 453 branches, from 151 genotypes (i.e. one branch from each of three replicate trees per genotype) in ten populations, representing all differentiated gene pools known in maritime pine (see Jaramillo-Correa *et al*., 2015). Every day between June 12^th^ and July 31^st^ 2018, one lateral branch per tree was cut from the previous year whorl in 30 of the selected trees (see above) and taken to the laboratory for inoculation. Inoculations were performed on the leader shoot of the current whorl of the excised branch.

### Experimental evaluation of susceptibility to Armillaria ostoyae

For the inoculation with *A. ostoyae*, we used the pathogen strain A4, collected from a dying maritime pine tree in La Teste (Gironde, France) in 2010 (Labbé *et al*., 2017b). For the experiment, two plugs of 5 mm diameter of malt agar with the *A. ostoyae* mycelium were put on the top of a mixture of industrial vegetable soup (Knorr 9 légumes©, Heilbronn, Germany), malted water and hazelnut wood chips in a 180 mL plastic jar (Heinzelmann & Rigling, 2016) (for a detailed laboratory protocol see Supporting Information S3.2). The lid was closed loosely enough to allow some oxygen flow. The jars were placed during three months in a heat chamber set at 23°C and 80% humidity before inoculation. The basal part of the sampled shoots (ca. 8 cm) was placed in the center of the mycelial culture in the heat chamber, maintaining the same temperature and humidity settings as for the mycelium growth, but adding an additional 12h cycle of light/dark. Only the jars showing a minimum jar occupation by *A. ostoyae* of 60% were used. After 3 weeks, wood colonization success for each sample was evaluated visually by confirming the presence of mycelium under the bark. At this point, the needles and the phloem of the branches were still green and thus the sample was considered adequate for susceptibility measurements. The length of the colonizing mycelium and length of the lesion in the sapwood (i.e. wood browning, hereafter referred to as necrosis) were measured. In the jar, we also visually evaluated the level of humidity of the medium (dry, medium, and very humid) and *A. ostoyae* growth. Controls were prepared in the exact same manner, but with plugs of sterile malt-agar as opposed to those colonized by *A. ostoyae*.

For this experiment, we randomly sampled ten maritime pine genotypes from six populations, one for each of the six differentiated gene pools represented in the CLONAPIN common garden. As root inoculations are destructive and impossible to implement directly in the common garden, we selected fully elongated current year shoots (bud stage 4 and 5), with a maximum diameter of 15 mm and a minimum length of 10 cm, as proxies (Matusick *et al*., 2016). A total of 180 branches from 60 genotypes (i.e. one branch from each of three replicate trees per genotype) were measured, cut and taken to the laboratory to be inoculated, on October 3^rd^-4^th^ 2018.

### Climatic Data

Summary climate data for the years 1950–2000 were retrieved for 32 variables from Worldclim (Hijmans *et al*., 2005) and a regional climatic model (Gonzalo, 2007) for the 11 non-Spanish and the 22 Spanish populations, respectively. Climate variables included monthly mean, highest and lowest temperatures, and mean monthly precipitation. Gonzalo’s (2007) model was favored for climate data in Spain because it considers a much denser network of meteorological stations than Worldclim, which is known to underperform, particularly for precipitation estimates, in this region (see Jaramillo-Correa *et al*., 2015).

### DNA extraction and SNP genotyping

Needles were collected from one replicate per genotype (*N*=416, including all genotypes used for pathogen susceptibility assays) and desiccated using silica gel. Genomic DNA was extracted using the Invisorb^®^ DNA Plant HTS 96 Kit/C kit (Invitek GmbH, Berlin, Germany). An Illumina Infinium SNP array developed by Plomion *et al*. (2016) was used for genotyping. Apart from potentially neutral genetic polymorphisms, this array comprises SNPs from candidate genes that showed signatures of natural selection (Eveno *et al*., 2008; Grivet *et al*., 2011), significant environmental associations with climate at the range-wide spatial scale (Jaramillo-Correa *et al*., 2015), or differential expression under biotic and abiotic stress in maritime pine (Plomion *et al*., 2016). After standard filtering followed by removal of SNPs with uncertain clustering patterns (visual inspection using *GenomeStudio v. 2.0)*, we kept 5,176 polymorphic SNPs, including 4,227 SNPs with a minor allele frequency (MAF) above 0.1.

### Quantitative genetic analyses

T o estimate the genetic variance components of the analyzed traits, we ran two different sets of mixed-effect models. The first one included three hierarchical levels, to account for the strong population genetic structure in maritime pine: gene pool, population nested within gene pool, and genotype (clone) nested within population and gene pool, as follows:

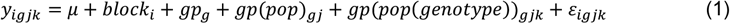

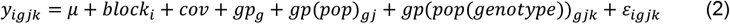

where for any trait *y_igjk_, μ* denotes the overall phenotypic mean, *block_i_* represents the fixed effect of experimental block *i, gp_g_* is the random effect of gene pool *g, gp*(*pop*)_*gj*_ is the random effect of pop *j* nested within gene pool *g, gp*(*pop*(*genotype*))_*gjk*_ is the random effect of genotype *k* nested within population *j* and gene pool *g*, and *ε* is the overall residual effect. In model 2, *cov* represents the covariates implemented when modeling the presence of pine processionary moth nests (i.e. tree height in 2015) and necrosis caused by *A. ostoyae* (i.e. a categorical evaluation of jar humidity). Using these models, the effects at the gene pool level had extremely wide confidence intervals, probably because some of them were represented by only a few highly contrasted populations. Thus, we also ran a set of models without the gene pool effect, as follows:

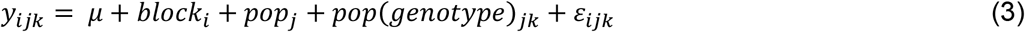

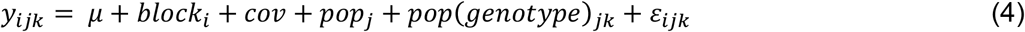

This set of models produced more accurate BLUP estimates, avoiding also confounded effects between gene pool and population, and was thus preferred for some of the subsequent analyses (see below).

All mixed-effect models were fitted in a Bayesian framework using Markov chain Monte Carlo (MCMC) methods implemented in the R package *MCMCglmm* (Hadfield, 2010) using R v.3.4.1 (R Development Core Team, 2017). All analyzed traits presented a Gaussian distribution with the exception of presence of pine processionary moth nests and needle discoloration caused by *D. sapinea* infection that followed a binomial distribution and were modeled with *logit* and *probit* link functions, respectively. Multivariate-normal prior distribution with mean centered around zero and large variance matrix (10^8^) were used for fixed effects with the exception of the model for needle discoloration caused by *D. sapinea* where a gelman prior for V was set, as suggested by Gelman *et al*. (2008) for ordinal traits. Inverse Wishart non-informative priors were used for the variances and covariances with a matrix parameter V set to 1 and a parameter n set to 0.002 (Hadfield, 2010). Parameter expanded priors were used to improve the convergence and mixing properties of the chain as suggested by Gelman (2006) for models on the presence of pine processionary moth nests, needle discoloration caused by *D. sapinea*, and necrosis caused by *A. ostoyae*. Parameter estimates were not sensitive to change in the priors. The models were run for at least 750,000 iterations, including a burn-in of 50,000 iterations and a thinning interval of 500 iterations. Four chains per model were run to test for parameter estimates convergence. Gelman-Rubin criterion Potential Scale Reduction Factor (psrf) was consistently below 1.01 (Gelman & Rubin, 1992) (see Table S4.1, Supporting Information, for further details on model specifications).

Variance components from the first set of models (including gene pool effect) were then used to compute broad-sense heritability (*H^2^*) as:

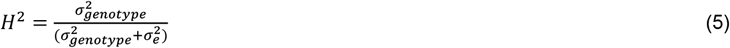

where 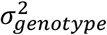 is the variance among genotypes within populations and gene pools and 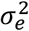 the residual variance. In addition, we computed the variance ratios associated to the population 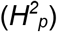 and gene pool 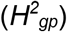 effects as:

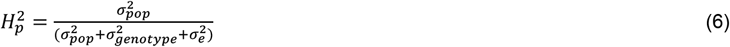

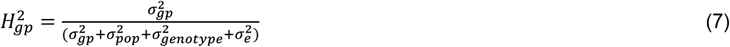

where 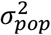 is the variance among populations within gene pools, 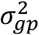 is the variance among gene pools. When appropriate, we included an extra term in the denominator to account for implicit *logit* and *probit* link function variance (π^2^/3 and +1, respectively; Nakagawa & Schielzeth, 2010). Genetic differentiation among populations for the analyzed traits (*Q*_ST_) was calculated as in Spitze (1993), using the second set of models (without gene pool effects, equations 3 and 4), as we were interested in the differentiation at the population level:

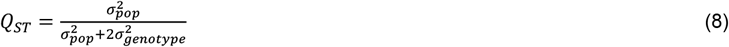

where 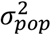 is the overall variance among populations and 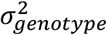 is the variance among genotypes within populations. Finally, we estimated the global *F*_ST_ using all available SNP genotypes in SPAGeDi 1.5 (Hardy & Vekemans, 2002). The difference between global *F*_ST_ and *Q*_ST_ values for each adaptive trait was considered significant when the 95% credible intervals (CI) did not overlap.

Genetic correlations were calculated with the Pearson’s coefficient of correlation using the Best Linear Unbiased Predictors (BLUPs) of the genotype (clone) effect (Henderson, 1973; Robinson, 1991) for each trait. The same estimate was used to characterize correlations of trait variation at population level considering the combined genotype and population effects. BLUPs were obtained from the model including genotype and population effects but not the gene pool effects (second set, equations 3 and 4), as they produced more precise estimates. In addition, we employed linear mixed models to test the effect of environmental variables on the population BLUPs while taking into account the assignment of each population to one of the six gene pools as random factor. First, we tested each model against a null-model without fixed effects to identify the significant associations. Then, for significant environmental associations, we calculated *R*^2^ as the proportion of variance explained by the fixed predictors using Nakagawa & Schielzeth’s approach (Johnson, 2014; Nakagawa & Schielzeth, 2013) implemented in the function ‘r2beta’ of the *r2glmm* package (Jaeger, 2017).

### Genotype-phenotype association

First, we used a mixed linear regression approach (MLM, Yu *et al*., 2006) implemented in Tassel v. 5.0 (Bradbury *et al*., 2007) to identify single SNPs associated to each of the phenotypes. Phenotypic values were estimated using the BLUPs accounting for both population and genotype effects from the models without gene pool effects (second set, equations 3 and 4). Ancestry proportions of each sample to six genetic clusters (see Jaramillo-Correa *et al*., 2015) were computed using STRUCTURE (Pritchard *et al*., 2000). These ancestry proportions were included as covariates in the MLM. A covariance matrix accounting for relatedness between all sample pairs was estimated using Loiselle’s kinship coefficient (Loiselle *et al*., 1995) in SPAGeDi 1.5 and was included as random effect. Negative kinship values were set to zero following Yu *et al*. (2006). Only loci with a *P*-value below 0.005 in the Tassel analyses and with a minimum allele frequency above 0.1 were used for further analyses. Second, for the identified associations, we used a Bayesian mixed-effect association approach (Bayesian Association with Missing Data, BAMD; Quesada *et al*., 2010; Li *et al*., 2012) in R to estimate single-locus genotype effects under three genetic models accounting for additive, over-dominance and dominance effects, respectively (as in Budde *et al*., 2014). As in the previous models, the STRUCTURE ancestry proportions were used as covariates and the relatedness matrix as random factor. Mean allelic effects (γ) and 95% confidence intervals were obtained from the distribution of the last 20,000 iterations (50,000 in total). Only those SNPs with credible intervals not overlapping zero were considered to have a significant (non-zero) effect on the trait.

Functional annotations, SNP motives and BLAST results for significant genotype-phenotype associations were retrieved from Plomion *et al*. (2016). Finally, in order to inspect visually geographical patterns, the minimum allele frequency of significantly associated SNPs was estimated in each population using SPAGeDi 1.5 and plotted in a map.

## Results

### Phenotypic variability, broad-sense heritability and genetic differentiation

Broad-sense heritability was highest for bud burst in 2015 (*H*^2^: 0.310, CI [0.272-0.358]), intermediate for height (*H*^2^: 0.219 [0.186-0.261]) and lowest for necrosis length caused by *A. ostoyae* (*H*^2^: 0.021 [0.004-0.121], Table 1). Broad-sense heritability of susceptibility to *D. sapinea*, assessed as the necrosis length, was also significant (*H*^2^: 0.152 [0.038-0.292]), with trees from northern Africa and southern Spain showing shorter necrosis length than trees from Atlantic populations (Figure 1a). Broad-sense heritability of needle discoloration caused by *D. sapinea* had similar magnitude, but was not significant (*H*^2^: 0.123 [0.000-0.233]). Necrosis length caused by *A. ostoyae* was most pronounced in southern populations, especially in Morocco and southern Spain, while less pathogen growth was observed in northern populations, e.g. in St. Jean de Monts from the French Atlantic gene pool (Figure 1b). Incidence of pine processionary moth nests in the common garden was not heritable (*H*^2^: 0.001 [0.000-0.207]). The tallest trees were found in populations from the Atlantic French and Atlantic Iberian gene pools, and for one of the Corsican populations (Pinia), whereas the shortest ones originated from southeastern Spain and Morocco (Figure S5.1, Supporting Information).

**Figure 1.**
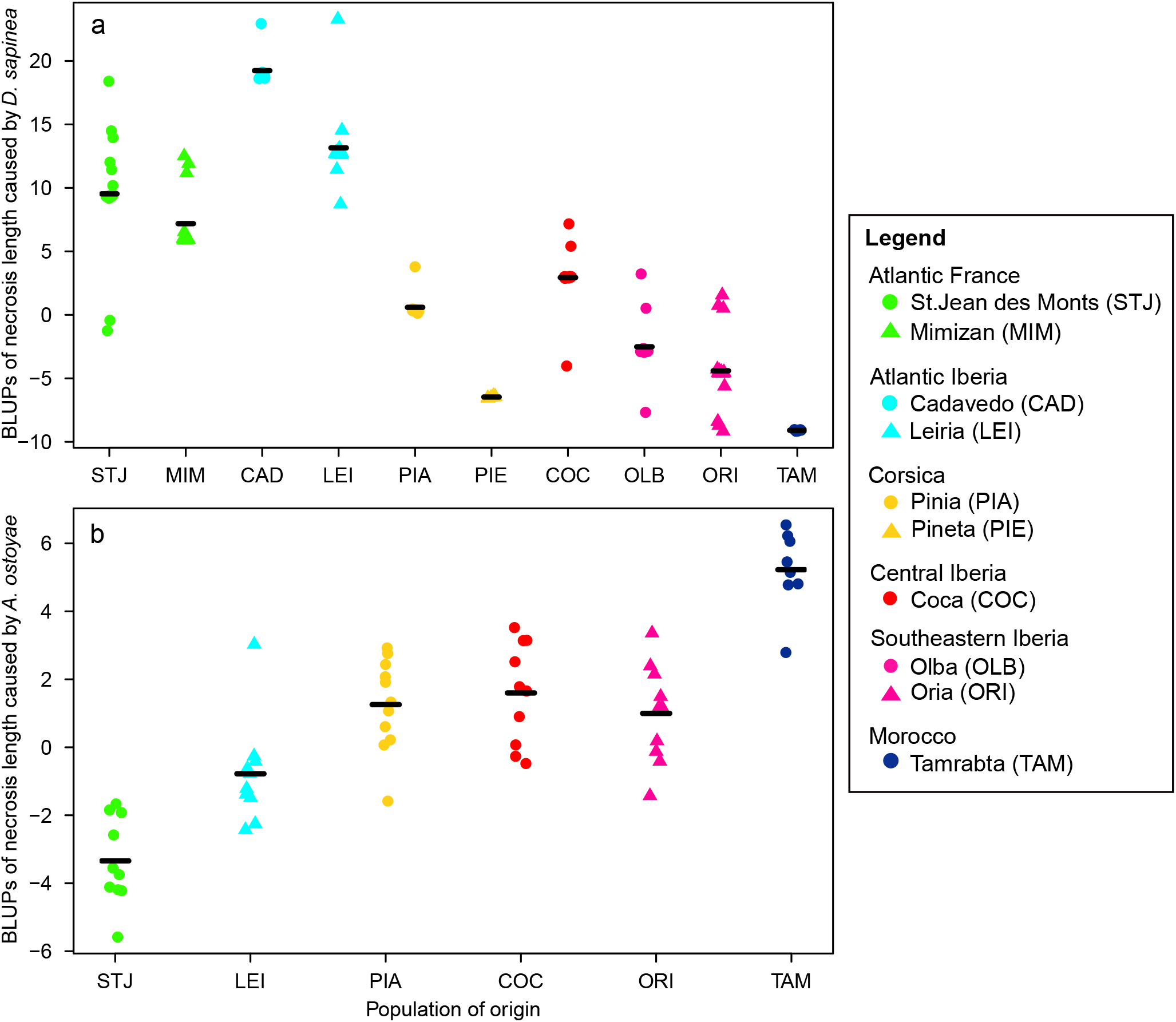
Stripchart of combined genotype and population BLUPs (Best Linear Unbiased Predictors) for necrosis length caused *by D. sapinea* (a) and *A. ostoyae* (b). Populations were assigned to one of the six gene pools identified by Jaramillo-Correa *et al*. (2015), corresponding to the six colours in the figure, and ordered by latitude (north to south). Symbols indicate different populations of the same gene pool. Black lines indicate the average necrosis length in each population.

**Table 1.**
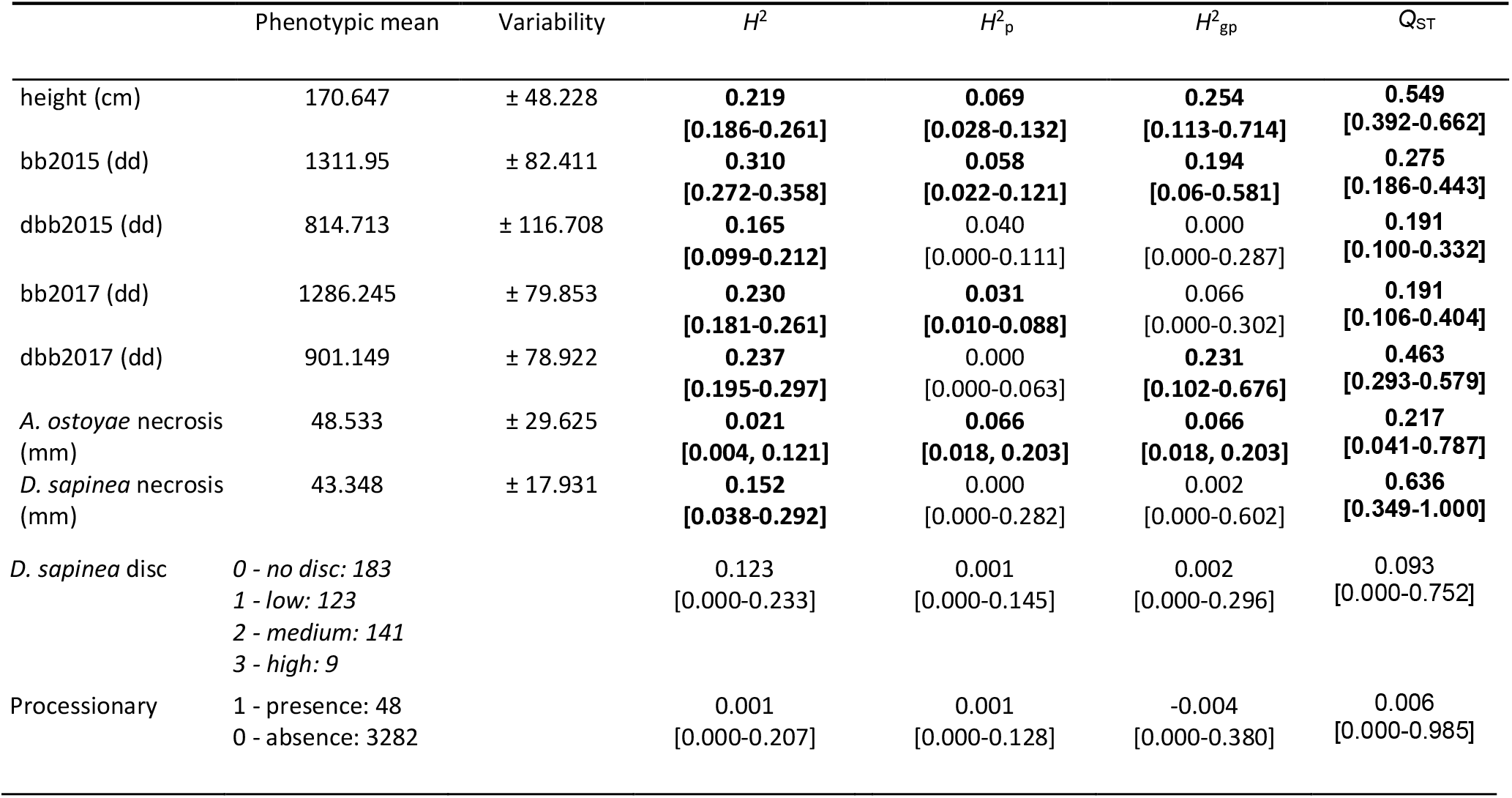
Broad-sense heritability, variance ratios and genetic differentiation of adaptive traits in *Pinus pinaster*. Variability refers to the standard deviation of the raw phenotypic data. *H*^2^, broad-sense heritability; *H*^2^_p_, variance ratio of the population effect (see main text for ratio definition); *H^2^_gp_*, variance ratio of the gene pool effect (see main text for ratio definition); *Q*_ST_, population differentiation at quantitative traits; bb, bud burst; dbb, duration of bud burst; disc, needle discoloration; processionary, presence/absence of processionary moth nests; dd, degree-days. Heritability and variance ratios for incidence of the processionary moth were computed using height as a covariate. For *A. ostoyae* necrosis length, the population and gene pool levels were identical as only one population per gene pool has been analyzed. Values in bold are significant. Values in squared brackets indicate the 95% credible intervals.

Most traits showed strong differences among gene pools, although the variance ratio associated to the gene pool effect was characterized by wide confidence intervals (Table 1). Based on overall population genetic differentiation, we observed significant *Q*_ST_ values (ranging from 0.191 for bud burst in 2017 and duration of bud burst in 2015, to 0.636 for necrosis length caused by *D. sapinea*) indicating strong population differentiation (Table 1). Global *F*_ST_ calculated using the available SNPs was 0.109 ([0.013; 0.325], p-value *Q*_ST_ estimates obtained for height and necrosis length caused by *D. sapinea* (Table 1).

### Correlations between traits and with environmental variables

All genetic correlations (i.e. those considering only the genotype effect) involving height and growth phenology were significant while disease susceptibility traits did not show significant genetic correlations except for necrosis length and needle discoloration caused by *D. sapinea* (Table 2). The strongest genetic correlation was observed between bud burst and duration of bud burst in 2015 (0.798, p-value < 0.001). Once the population effect was considered, a significant negative trait covariation was found between necrosis length caused by each of the two fungal pathogens (−0.692, p-value < 0.001; Table 2, Figure 2). We also observed significant population correlations with height, negative for necrosis length caused by *A. ostoyae* (−0.653, p-value < 0.001) and positive for necrosis length caused by *D. sapinea* (0.679, p-value < 0.001).

**Figure 2.**
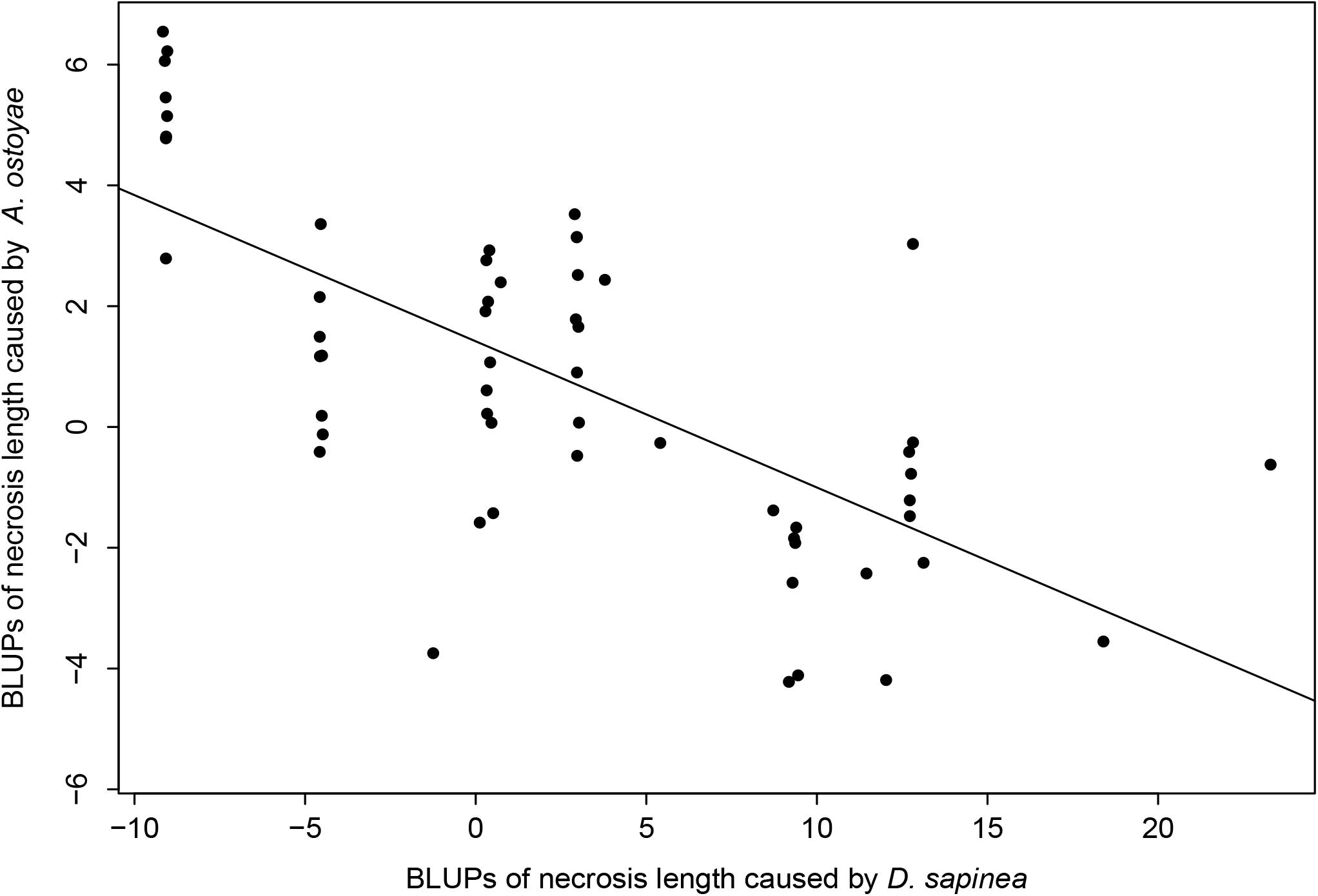
Correlation of trait variation across *Pinus pinaster* populations based on BLUPs (Best Linear Unbiased Predictors) for necrosis length caused by *Diplodia sapinea* and *Armillaria ostoyae*. A linear trend line is also shown (Pearson’s correlation coefficient = - 0.692, p-value<0.001).

**Table 2.** Pearson’s correlation coefficients of the Best Linear Unbiased Predictors (BLUPs) of genotype (i.e. genetic correlations, above diagonal) and combined population and genotype effects (below diagonal). bb, bud burst; dbb, duration of bud burst; disc, needle discoloration. Significance levels after false discovery rate (FDR) correction: *<0.05; **<0.01; ***0.001.

|  | height | bb2015 | dbb2015 | bb2017 | dbb2017 | *D. sapinea* necrosis | *D. sapinea* disc | *A. ostoyae* necrosis |
| --- | --- | --- | --- | --- | --- | --- | --- | --- |
| height |  | -0.172** | -0.132* | -0.178** | 0.439*** | 0.111 | 0.082 | -0.010 |
| bb2015 | 0.194*** |  | 0.798*** | 0.392*** | -0.292*** | 0.118 | 0.041 | -0.017 |
| dbb2015 | 0.117* | 0.846*** |  | 0.380*** | -0.233*** | 0.061 | 0.065 | -0.110 |
| bb2017 | 0.021 | 0.593*** | 0.547*** |  | -0.154** | -0.023 | 0.069 | -0.092 |
| dbb2017 | 0.800*** | 0.147** | 0.079 | 0.018 |  | -0.032 | -0.093 | -0.030 |
| *D. sapinea* necrosis | 0.679*** | 0.564*** | 0.511*** | 0.309*** | 0.639*** |  | 0.231* | -0.009 |
| *D. sapinea* disc | 0.414*** | 0.470*** | 0.492*** | 0.337*** | 0.392*** | 0.727*** |  | -0.149 |
| *A. ostoyae* necrosis | -0.653*** | -0.321* | -0.284* | -0.056 | -0.530*** | -0.692*** | -0.367** |  |

Moreover, using mixed linear models that account for gene pool as random factor, we observed a negative relationship for maximum temperatures in the summer months with height, duration of bud burst in 2017, and necrosis length and needle discoloration caused by *D. sapinea* (Table 3). The strongest relationship was found for height and maximum temperatures in July (*R^2^* = 0.709 [0.556-0.831], p-value <0.001) while the variance of e.g. needle discoloration caused by *D. sapinea* explained by maximum temperatures in July was lower (*R*^2^ = 0.524 [0.128-0.841], p-value = 0.004; Table 3, Figure 3). Also, geographical variables, such as altitude, latitude and longitude showed significant correlation with all phenotypes. Interestingly, necrosis length caused by *A. ostoyae* showed a latitudinal cline (*R*^2^ = 0.750, p-value = 0.016) but only a marginally significant positive correlation with maximum temperatures in July (*R*^2^=0.514, p-value= 0.066).

**Figure 3.**
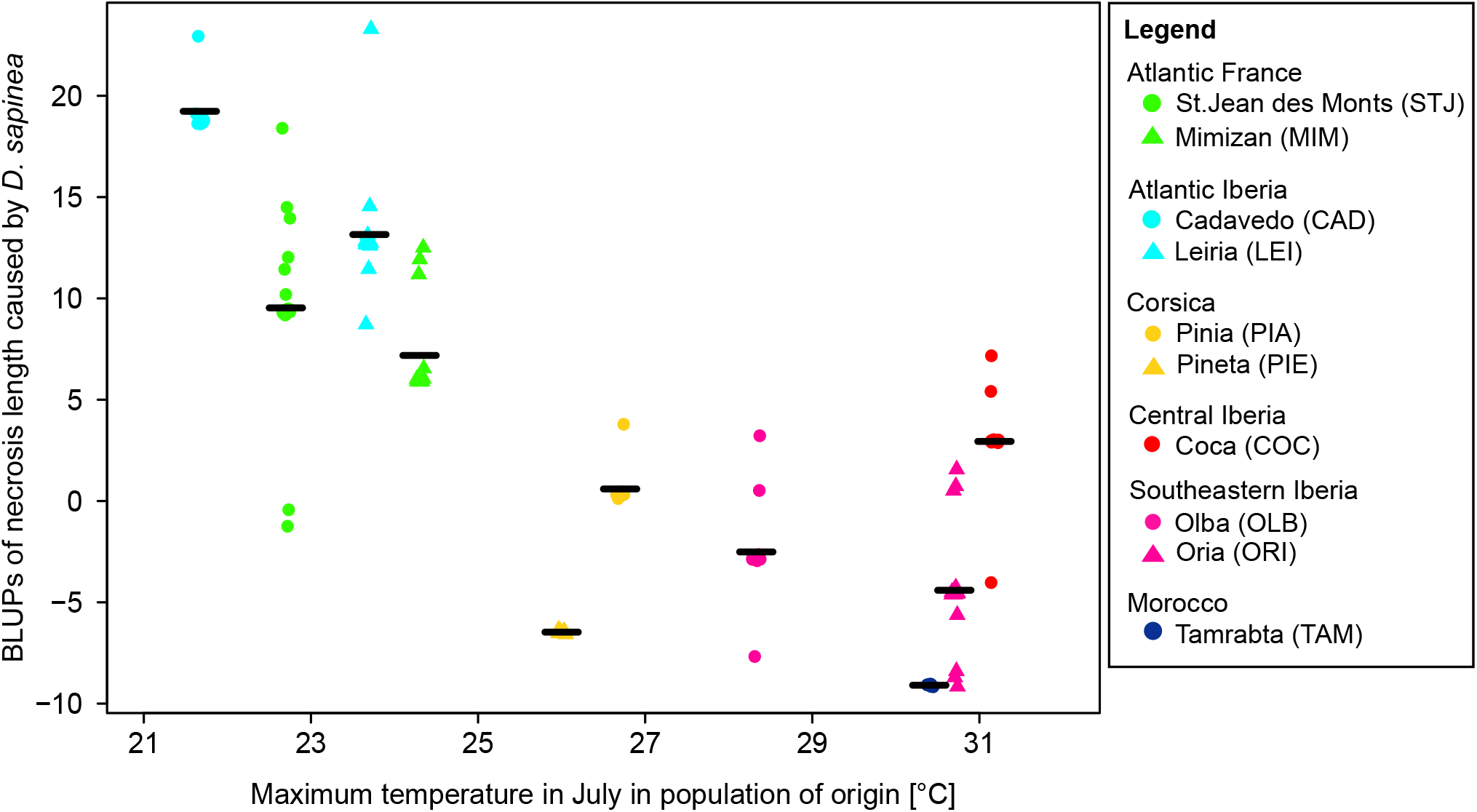
Stripchart of combined genotype and population BLUPs (Best Linear Unbiased Predictors) for necrosis length caused *by D. sapinea* plotted against the maximum temperature in July in the population of origin. Populations were assigned to one of the six gene pools identified by Jaramillo-Correa *et al*. (2015), corresponding to the six colours in the figure. Symbols indicate different populations of the same gene pool. Black lines indicate the average necrosis length in each population.

**Table 3.** Correlations between adaptive phenotypes and environmental variables, as identified using linear mixed-effects models that account for gene pool as a random factor. Only models that performed significantly better than a null model excluding the fixed effect are reported. *R*^2^, the proportion of variance explained by the fixed predictor in the mixed-effects model. For necrosis length caused by *A. ostoyae*, a simple linear regression model was used as each gene pool was only represented by a single population. bb, bud burst; dbb, duration of bud burst; disc, needle discoloration; Tmax, maximum temperature; prec, precipitation.

| Trait | Environmental factor | $R^2$ | $P$ |
| --- | --- | --- | --- |
| height | altitude | 0.650 | <0.001 |
|  | Tmax July | 0.709 | <0.001 |
|  | Tmax August | 0.201 | 0.023 |
| bb2015 | altitude | 0.200 | 0.020 |
| dbb2015 | longitude | 0.245 | 0.044 |
| bb2017 | longitude | 0.262 | 0.028 |
| dbb2017 | latitude | 0.290 | 0.042 |
|  | altitude | 0.430 | 0.001 |
|  | Tmax July | 0.212 | 0.044 |
|  | Tmax August | 0.193 | 0.033 |
|  | prec in August | 0.123 | 0.027 |
|  | prec in September | 0.123 | 0.027 |
| *D. sapinea* necrosis | latitude | 0.341 | 0.031 |
|  | altitude | 0.482 | 0.018 |
|  | Tmax July | 0.465 | 0.034 |
|  | Tmax August | 0.394 | 0.036 |
| *D. sapinea* disc | latitude | 0.284 | 0.013 |
|  | Tmax June | 0.310 | 0.009 |
|  | Tmax July | 0.524 | 0.004 |
|  | Tmax August | 0.462 | 0.003 |
|  | Tmax September | 0.315 | 0.004 |
| *A. ostoyae* necrosis | latitude | 0.750 | 0.016 |
|  | altitude | 0.614 | 0.040 |

### Genotype-phenotype association

Between three and 28 SNPs were significantly associated with each of the phenotypic traits evaluated under different genotype effect models (see Table S6.1, Supporting Information). Here we only focus on SNPs that were significant under the additive genetic model, this model being built on three genotypic classes and therefore considered the most robust. Based on this model, five SNPs were associated to height, 27 SNPs were associated with growth phenology (considering altogether the different phenology traits and measurement years), and seven with pathogen susceptibility (Table 4). In total, four significantly associated SNPs showed non-synonymous changes. Two non-synonymous SNPs were associated to bud burst in 2017 (Figure S6.1, Supporting Information). In addition, one non-synonymous SNP was associated to needle discoloration caused by *D. sapinea* and another one to duration of bud burst in 2015 (Table 5 and Figure 4). All the remaining SNPs involved in significant associations under the additive model were either non-coding or the effect of the substitution was unknown (Table S6.1, Supporting Information). The geographical distribution of minor allele frequency of the associated SNPs was quite variable and did not reflect the population genetic structure of the species (Figure 5).

**Figure 4.**
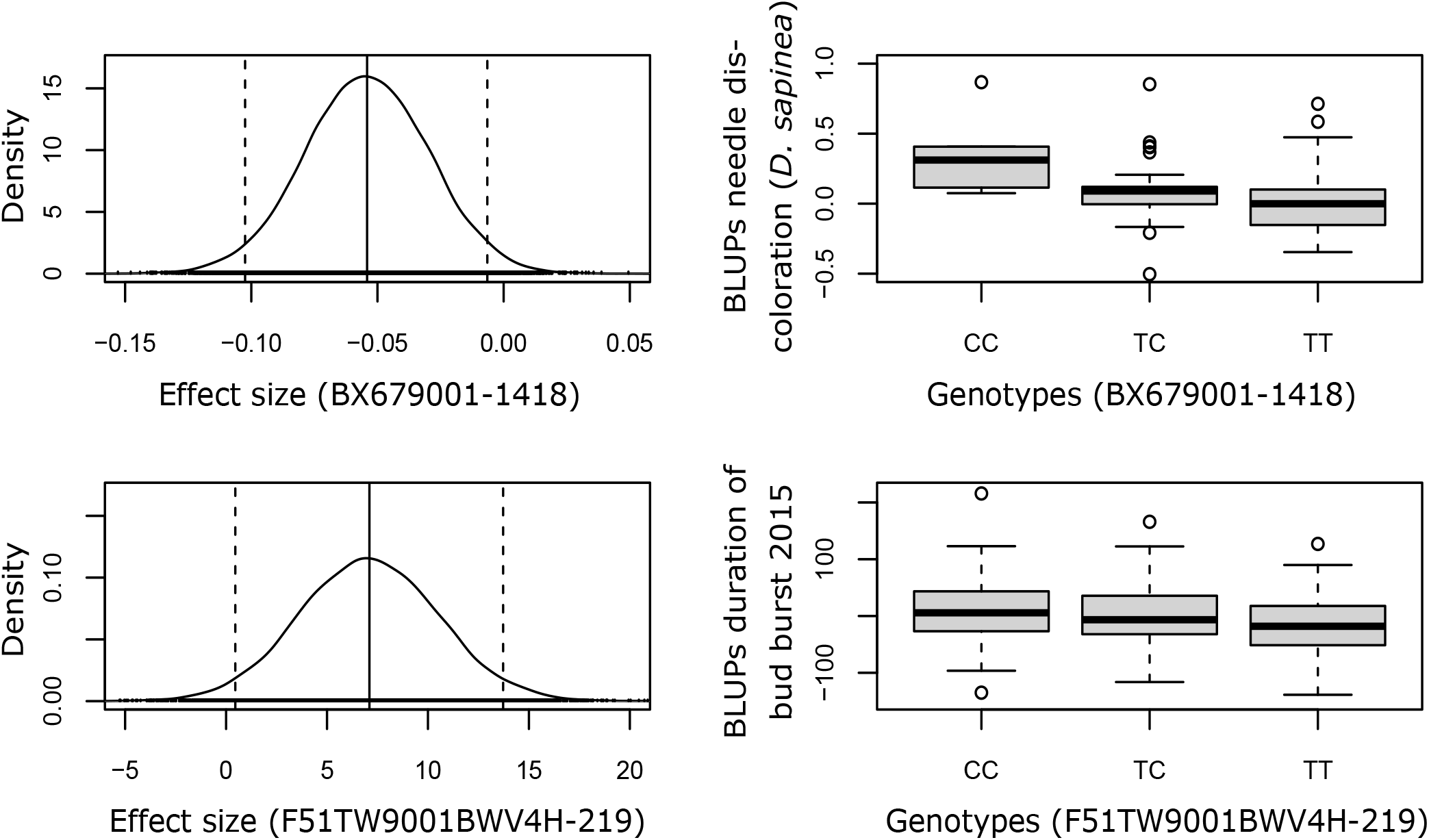
Density plots of the effect sizes based on 20,000 BAMD simulations (left) and genotypic effects (box plots, right) for two exemplary Single Nucleotide Polymorphisms (SNPs) showing significant association with needle discoloration caused by *Diplodia sapinea* (above) and duration of bud burst in 2015 (below) in *Pinus pinaster*.

**Figure 5.**
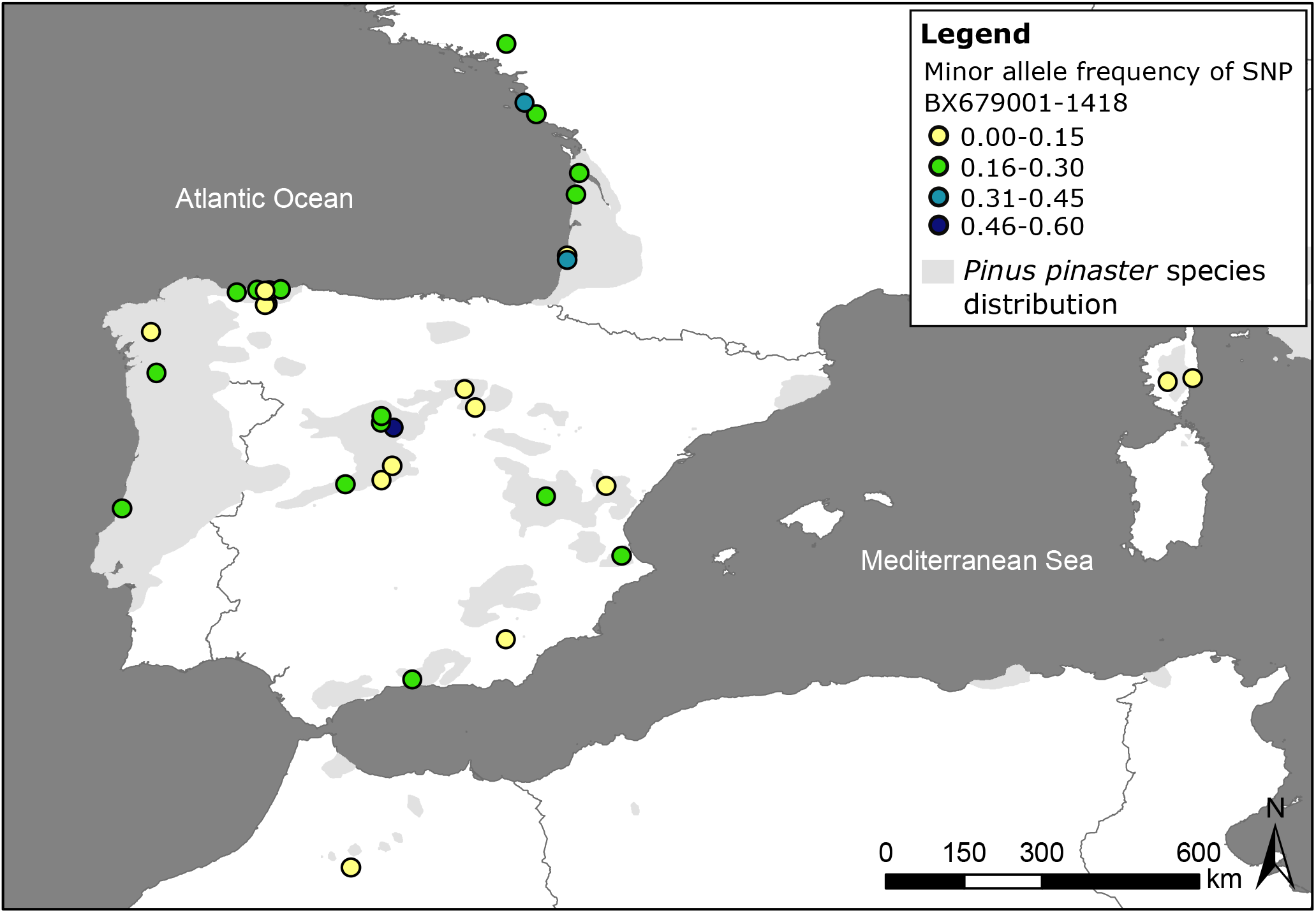
Minor allele frequency distribution of SNP BX679001-1418 in natural populations of *Pinus pinaster*. This locus was significantly associated to needle discoloration caused by *Diplodia sapinea*, and possibly coded for a disease-induced translation initiation factor, eIF-5.

**Table 4.**
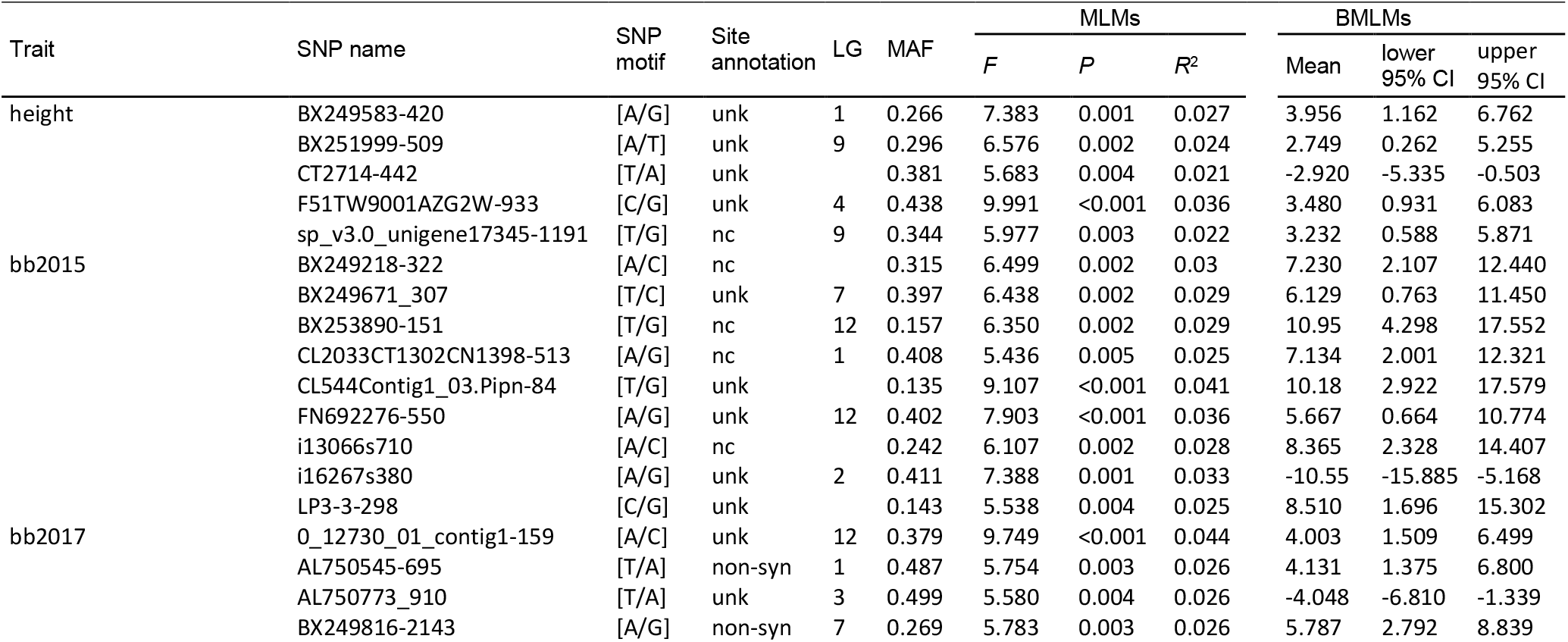
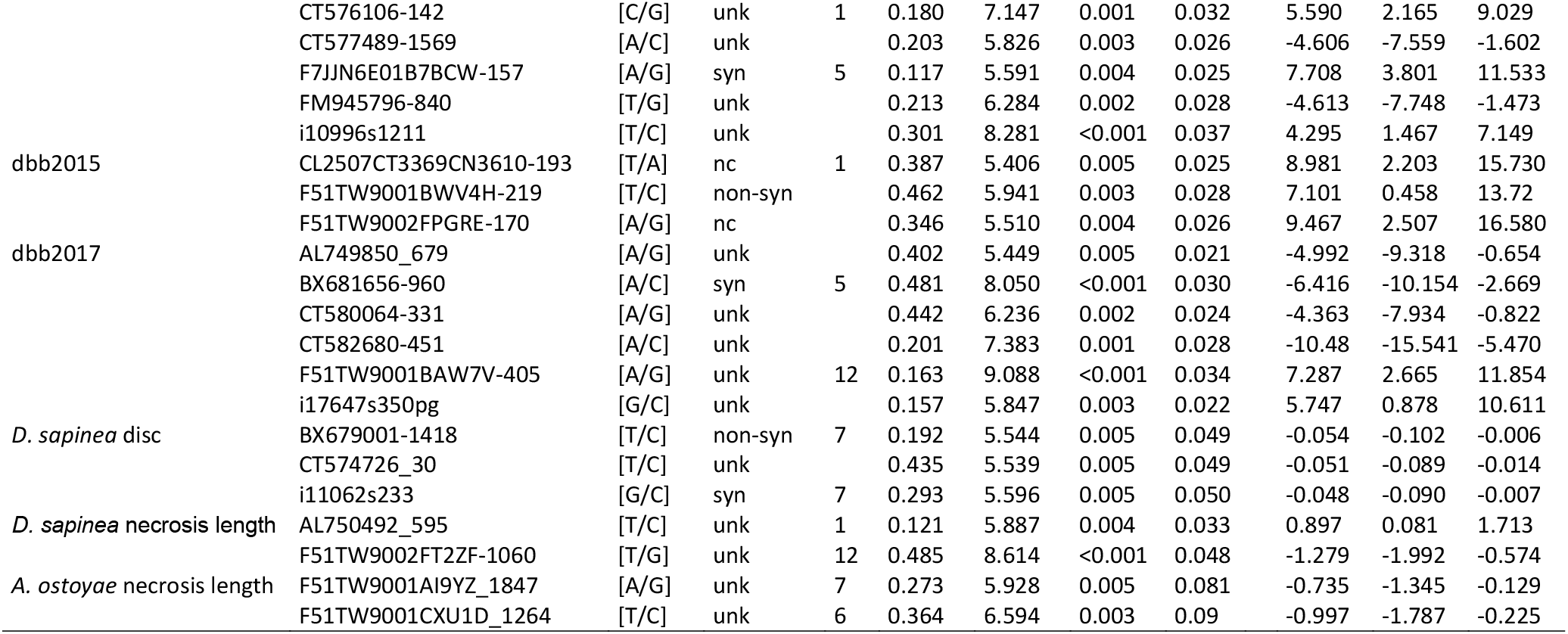
Single Nucleotide Polymorphisms (SNPs) significantly associated to height, growth phenology and pathogen susceptibility traits under the additive genetic model, as identified by a two-step approach based on mixed-effects linear models (MLMs) implemented in Tassel and the Bayesian framework in BAMD (BMLMs). Bayesian mean SNP-effects and 95% credible intervals (CIs) were obtained from the distribution of the last 20,000 iterations in BAMD. Marker codes and linkage groups as reported in Plomion *et al*. (2016). bb, bud burst; dbb, duration of bud burst; disc, needle discoloration; LG, linkage group; MAF, minimum allele frequency; unk, unknown; nc, noncoding; non-syn, non-synonymous; syn, synonymous.

**Table 5.** Annotation for SNPs significantly associated with adaptive traits under the additive model and coding for a non-synonymous amino acid change, as retrieved from Plomion *et al*. (2016). bb, bud burst; dbb, duration of bud burst; disc, needle discoloration.

| Trait | SNP name | Motif | Protein change | Putative protein function |
| --- | --- | --- | --- | --- |
| dbb2015 | F51TW9001BWV4H-219 | [T/C] | Asparagine - Serine | LANC-like domain containing protein |
| bb2017 | BX249816-2143 | [A/G] | Glutamate/Glutamine - Arginine | putative Cdc2-related protein kinase<br>CRK2 |
| bb2017 | AL750545-695 | [A/T] | Glutamate/Glutamine - Valine | Catalase |
| *D. sapinea* disc. | BX679001-1418 | [T/C] | Isoleucine - Valine | Translation initiation factor eIF-5 |

## Discussion

In the current context of climate change, understanding the genetic basis of adaptive traits in tree species is key for an informed forest management and the development of new breeding applications. In this study, we assessed variation in maritime pine for height and growth phenology traits, as well as for incidence of pine processionary moth nests and response to two pathogenic fungi, *D. sapinea* and *A. ostoyae*, at the range-wide scale. Novel inoculation protocols were developed for *D. sapinea* and *A. ostoyae* based on excised branches and applied to trees grown in a clonal common garden. For the first time, we estimated broad-sense heritability of pine susceptibility for these two major fungal pathogens (evaluated as necrosis length), as well as the variance ratios associated to the population and gene pool effects. Most of these adaptive traits showed significant broad-sense heritability. Variation of pathogen susceptibility among geographical provenances, as well as duration of bud burst in 2017, followed a latitudinal gradient, possibly corresponding to a climatic gradient, but in opposite directions for the two pathogens. A genetic association approach revealed several loci significantly associated to height, growth phenology and pathogen susceptibility in maritime pine. This information will be useful for current efforts to implement breeding strategies based on genome prediction (i.e. genomic selection) in this species. The presence of pine processionary moth nests evaluated in the common garden was not heritable but future studies should consider the level of infestation and/or assess damage more precisely.

### Genetic variation and differentiation, and correlations with climate

Broad-sense heritability estimates were in the range of previously published values in maritime pine or other forest tree species. The variance ratios associated to population and gene pool effects reflect genetic differences that are not necessarily due to selection but might also reflect other evolutionary processes at the population level, e.g. drift or different levels of gene flow. Nevertheless, they provide important insights on trait variation at the range-wide geographical scale. The variance ratio associated to the gene pool effect had very wide confidence intervals indicating that highly differentiated units were mixed at this level. Already Rodríguez-Quilón *et al*. (2016) described strong quantitative genetic differentiation for height and survival within gene pools of maritime pine. While gene pools may reflect different evolutionary histories, our results suggest that selection and hence adaptation at quantitative traits takes place at a finer spatial scale. Therefore, genetic differentiation for adaptive traits at the population level (e.g. as estimated by *Q*_ST_) is expected to be more informative. Neutral genetic differentiation, i.e. *F*_ST_, was moderate (*F*_ST_ = 0.109 [0.013; 0.325], p-value < 0.001) and significantly lower than *Q*_ST_ estimates obtained for height and necrosis length caused by *D. sapinea*, which suggests that divergent selection is promoting local adaptation in these traits (Whitlock & Guillaume, 2009; Lamy *et al*., 2011).

Height is a crucial, frequently studied trait in forest trees, as increased height growth is a main target of breeding programmes (e.g. Kremer & Lascoux, 1988; Cornelius, 1994). In our study, we found moderate broad-sense heritability of 0.219 for height, well in line with estimates in other conifer species e.g. ranging from 0.21 in *Pinus taeda* to 0.78 in *Picea abies* (reviewed in Lind *et al*., 2018) and from 0.148 to 0.282 in maritime pine saplings depending on the common garden site and the provenance (Rodríguez-Quilón *et al*., 2016). Height is known to be a highly integrative trait closely responding to abiotic factors, such as climate (Alía *et al*., 2014; Jaramillo-Correa *et al*., 2015), and has thus been used in combination with genetic markers to identify relevant conservation units in maritime pine (Rodríguez-Quilón *et al*., 2016). In our study, we showed that height is not only correlated with climate but also with biotic factors, such as pathogen susceptibility (positively for *D. sapinea* and negatively for *A. ostoyae)*. This is crucial information, for example, for the Landes maritime pine breeding programme, which is based on artificial selection for height growth and straightness, and that may have selected at the same time for higher susceptibility to *D. sapinea*, a pathogen that is expected to increase its presence in maritime pine productive forests in the near future (see below).

Growth phenology traits (date and duration of bud burst) showed low to moderate broadsense heritability, depending on the year (2015 or 2017). Differentiation (*Q*_ST_) for bud burst varied from 0.191 to 0.275, which is comparable to a mean of 0.249 for bud flush averaged over several forest tree species (reviewed in Alberto *et al*., 2013). In our study, trees originating from northern populations flushed later than trees from southern populations. Similar clines have been observed for other conifers (reviewed in Alberto *et al*., 2013), which is not surprising, as spring phenology, such as flushing time, is known to be correlated with climatic factors (e.g. Zohner & Renner, 2014). Such clines in growth phenology along climate gradients suggested local adaptation due to divergent selection, which might suffer mismatch with changing climate conditions (Badeck *et al*., 2004; Lindner *et al*., 2010, Alberto *et al*., 2013). Spring phenology can also play a role in resistance to or avoidance of forest tree pathogens (e.g. Swedjemark *et al*., 1998; Ghelardini & Santini, 2009; Nielsen *et al*., 2017). In line with this, we found a positive genetic correlation between needle discoloration and necrosis length caused by *D. sapinea* with growth phenology indicating that earlier flushing trees with faster spring growth showed less severe disease symptoms. Krokene *et al*. (2012) showed that the concentrations of starch and total sugars (glucose, fructose and sucrose) in twigs of *Picea abies* change during shoot development, which affects pathogen-related symptoms. In our study, inoculations were carried out on twigs with already elongated needles, however, the chemical composition of twigs might differ with time since bud burst and we cannot fully discard these effects in our assessment of disease incidence.

Broad-sense heritability for susceptibility to *D. sapinea* based on necrosis length (*H^2^*=0.152) was much lower than e.g. narrow-sense heritability of lesion length caused by *Fusarium circinatum* (*h^2^*=0.618, Elvira-Recuenco *et al*., 2014). However, this low but significant heritability still indicated the potential for an evolutionary response of maritime pine to *D. sapinea*. Moreover, *Q*_ST_ was highest for necrosis length caused by this pathogen (*Q*_ST_=0.636) in our study, which was also significantly higher than neutral-marker *F*_ST_. This result pointed to adaptive divergence and substantial variability in susceptibility to *D. sapinea* across the geographical range of maritime pine. This population variability could be harnessed in maritime pine breeding programmes to increase resistance to the disease (Bouffier *et al*., 2008, Ingvarsson & Dahlberg, 2019). Remarkably, we observed a negative correlation of trait variation across populations for necrosis length caused by *D. sapinea* and *A. ostoyae*. This direct comparison should be taken with caution as the inoculation experiments were carried out at different time points (*D. sapinea* in summer and *A. ostoyae* in autumn), which is likely to affect the host response. However, correlation with climate across populations for susceptibility to both *D. sapinea* and *A. ostoyae* (as well as for other adaptive traits such as height and growth phenology) indicated similar environmental clines driving differentiation at these traits. Especially, maximum temperatures during summer months showed significant correlations with genetic variability of susceptibility to pathogens across maritime pine populations. Trees from populations with high maximum summer temperatures were less susceptible to *D. sapinea*. This result can be interpreted in different ways: 1) If we assume that *D. sapinea* is native in Europe, the pathogen pressure can be expected to be stronger in southern regions, with a climate more favourable to *D. sapinea* pathogenic outbreaks, triggered by stress in the host plant, especially by droughts (Luchi *et al*., 2014). Maritime pine populations growing in these regions (e.g. Morocco and southern Spain) would then be more likely to have evolved resistance to the disease. In contrast, trees from populations where severe drought periods have most likely not been common so far (e.g. Atlantic populations from the Iberian Peninsula and France) would be more susceptible. 2) If maritime pine and *D. sapinea* have not had sufficient time to co-evolve (e.g. if *D. sapinea* was not native to Europe) or pathogen pressure was not strong enough, differences in susceptibility among maritime pine populations might be due to exaptation or ecological fitting, i.e. due to correlated traits selected for other functions (Agosta & Klemens, 2008). Populations of maritime pine strongly vary geographically in many traits related to growth and response to drought, along the gradient from North Africa to the Atlantic regions of the Iberian Peninsula and France (Correia *et al*., 2008; Aranda *et al*., 2010; Corcuera *et al*., 2012; Gaspar *et al*., 2013; de la Mata *et al*., 2014). Some of these traits may indirectly influence their susceptibility to pathogens, as observed here for *D. sapinea*. For example, faster growing maritime pine trees from northern populations are known to invest more in inducible defences while slow growing trees from southern populations invest more in constitutive defences (López-Goldar *et al*., 2018). The positive correlation between height and necrosis length caused by *D. sapinea* might indicate that constitutive defences confer better resistance to this pathogen in the southern populations. Also, Meijón *et al*. (2016) showed that the metabolomes in needles of maritime pine trees from populations with distinct geographic origin (notably Atlantic versus Mediterranean provenances) were quite differentiated, with flavonoids showing a significant correlation with the water regime of the population of origin. However, the expression of metabolites is organ specific (de Miguel *et al*., 2016) and knowledge about secondary metabolites involved in resistance to *D. sapinea* is still lacking.

A study on the invasive pathogen *Fusarium circinatum*, which did certainly not co-evolve with maritime pine, also revealed a geographic cline in susceptibility, with Atlantic maritime pine populations showing less susceptibility than Moroccan populations (Elvira-Recuenco *et al*., 2014). A similar pattern was observed for *A. ostoyae* in our study. Our results indicated that maritime pine from southwestern France, where *A. ostoyae* outbreaks have been frequently reported (Labbé *et al*., 2015), may have developed some resistance or might show exapted resistance to the disease. Considering the absence of reports of *A. ostoyae* from the south of the Iberian Peninsula (Marxmüller & Guillaumin, 2005), which is in line with the species’ preference for humid forest sites (Cruickshank *et al*., 1997; Heinzelmann *et al*., 2019), trees in Morocco and southern Spain have most-likely never co-evolved with this pathogen. However, a study by Guillaumin *et al*. (2005) on the mortality of potted maritime pine plants revealed an opposite pattern to the one found in our study, with the Landes population in Atlantic France being the most susceptible and the Moroccan population the least susceptible to *A. ostoyae*. Moreover, Zas *et al*. (2007) found much higher (narrow-sense) heritability (*h*^2^=0.35) for mortality due to *A. ostoyae* in an infested progeny trial of maritime pine seedlings than the values of broad-sense heritability of necrosis length found in our study. *Armillaria ostoyae* is a root pathogen and a critical point during natural infestation that could be key for resistance mechanisms is the penetration into the root (Prospero *et al*., 2004; Solla *et al*., 2011; Labbé *et al*., 2017b), as the pathogen grows faster once it enters the organism and reaches the cambium (Solla *et al*., 2002). This step was bypassed in our inoculation protocol on excised branches. In the future, it would therefore be interesting to carry out inoculations on potted seedlings or young trees from range-wide maritime pine populations to confirm the patterns observed in this study.

Suitable strategies to evaluate susceptibility to *D. sapinea* and *A. ostoyae* will become increasingly important as climate change increases pathogen pressure. Droughts are expected to become more frequent throughout Europe (IPCC, 2014), which will most likely trigger *D. sapinea* outbreaks also in regions where the pathogen has not caused severe disease symptoms so far. Recently, a northward expansion of *D. sapinea* outbreaks in Europe, probably driven by higher spring temperatures, has been reported; these outbreaks have caused severe damage on *P. sylvestris* in Sweden and eastern Baltic countries (Adamson *et al*., 2015; Brodde *et al*., 2019). Because some of the largest and most productive maritime pine forests are located in the Atlantic distribution of the species (e.g. the Landes region in southwestern France), our results suggest that the expected increase of drought events in these populations will most likely cause severe damages due to their high susceptibility to *D. sapinea*. In the case of *A. ostoyae*, the main threat resides in the condition of the host. As mentioned before, a weaker host will be more susceptible to the fungus, and future extreme weather events are bound to weaken trees, also increasing pathogenicity of *A. ostoyae* (Kubiak *et al*., 2017). A mathematical model predicted a drastic northward shift of *A. ostoyae* in the Northwestern United States for the years 2061-2080, leading to increased mortality of stressed and maladapted trees (Hanna *et al*., 2016). As suggested by this study, trees maladapted to new temperatures are also expected to be more susceptible to biotic stress.

### Genotype-phenotype association and candidate genes

By using a single-locus, two-steps approach, we revealed significantly associated loci for all heritable traits under study. However, genotype effects were small, pointing to a highly polygenic nature of the studied traits, as often reported for adaptive traits in forest trees. In addition, for susceptibility to *D. sapinea* and *A. ostoyae*, no resistance alleles with major effects were detected. Nevertheless, information on genotype-phenotype associations can make genome-based phenotypic predictions more efficient by providing a set of SNPs that are more closely linked to causal SNPs than those selected just based on genome location (e.g. Westbrook *et al*., 2013). We retrieved annotations from Plomion *et al*. (2016) and found four non-synonymous SNPs significantly associated to duration of bud burst in 2015 (one locus), bud burst in 2017 (two loci) and needle discoloration caused by *D. sapinea* (one locus) under the additive genetic model (see Table 5). The potential function of these genes has to be interpreted with caution as this information usually derives from studies in distantly related model species (typically *Arabidopsis thaliana*). Nevertheless, the locus (BX679001-1418), which was significantly associated to needle discoloration caused by *D. sapinea*, possibly codes for a translation initiation factor, eIF-5, that has previously been reported to be involved in pathogen-induced cell death and development of disease symptoms in *A. thaliana* (Hopkins *et al*., 2008). This gene deserves further attention in future studies addressing the genetic control of adaptive traits in conifers. Also, differential gene expression analyses could shed light on the genes and pathways involved in pathogen resistance in maritime pine, when applied on inoculated and control trees growing under controlled conditions in greenhouses or common gardens.

Based on a well-replicated clonal common garden and state-of-the-art genotyping technology, we were able to study key adaptive traits in maritime pine and found evidence for non-synonymous mutations underlying genetic variation for some of these traits. Association studies for highly polygenic traits are still challenging. Lind *et al*. (2017) reported an average of 236 SNPs (out of 116,231 tested, i.e. 0.203 %) associated to each of four fitness-related traits in *Pinus albicaulis* by detecting signals of significantly higher covariance of allele frequencies. In the near future, multilocus association methods should be used to reveal genome wide loci with non-zero effects for polygenic traits in forest trees (Goldfarb *et al*., 2013; De la Torre *et al*., 2019).

### Conclusions

In our study, we evaluated phenotypic variability and broad-sense heritability for height, growth phenology, pathogen susceptibility and incidence of processionary moth nests in a range-wide clonal common garden of maritime pine. We revealed strong genetic divergence at several adaptive traits, especially height and necrosis length caused by *D. sapinea*, across maritime pine populations. We have also shown that several adaptive traits in maritime pine were genetically correlated and that population variation was often established along climatic clines, in particular with maximum temperatures. The evolution of suits of functional traits along environmental clines is a common pattern in nature (e.g. Chapin *et al*., 1993; Reich *et al*., 1996). Currently, locally adapted populations are challenged by changing climate conditions, and emergent pests and pathogens expanding their range (Seidl *et al*., 2017). Susceptibility to *D. sapinea* was highest in the Atlantic maritime pine populations where it is expected to cause severe outbreaks due to increased incidence of drought events in the future. Correlated selection for increased susceptibility needs to be considered in breeding programmes aiming at increasing height growth in this species. In addition, opposing trends in pathogen susceptibility across maritime pine populations, e.g. for *D. sapinea* and *A. ostoyae* (this study), and for the invasive pathogen *F. circinatum* (Elvira-Recuenco *et al*., 2014), challenge forest tree breeding and natural forest resilience. An improved understanding of integrated phenotypes, including responses to known pests and pathogens, and their underlying genetic architecture is fundamental to assist new-generation tree breeding and the conservation of valuable genotypes. Coupled with early detection methods (see e.g. Kenis *et al*., 2018), knowledge on genetic responses to emerging pests and pathogens will help to ensure the health of forests in the future. Finally, given the recent development of efficient technologies to combine functional genetic variants (e.g. genome editing), the identification of large numbers of polymorphisms associated with commercial traits is expected to contribute to a new era in plant breeding (Breeding 4.0, see Wallace *et al*., 2018).

## Acknowledgements

We thank Xavier Capdevielle, Olivier Fabreguettes, Martine Martin-Clotté and Gilles St Jean for field and lab assistance, and Brigitte Lung-Escarmant, Thierry Belouard, Bernard Boutte, Claude Husson, Jean-Baptiste Daubrée, Margarita Elvira-Recuenco and Rosa Raposo-Llobet for valuable discussions. We thank Juan Majada for providing the rooted cuttings used to establish the CLONAPIN collection in Cestas and the experimental unit of INRA-Pierroton for trial establishment, height and bud flush measurements. We thank Hervé Jactel and Victor Rebillard for providing the processionary moth nest data. We acknowledge funding from IdEx Bordeaux - Chaires d’installation 2015 (EcoGenPin), the Spanish National Research Plan (ClonaPin, RTA2010-00120-C02-01), and the European Union’s Horizon 2020 research and innovation program under grant agreement No 773383 (B4EST). This work is part of the PhD research of A.H. funded by the Région de Nouvelle-Aquitaine (project Athénée) and the IdEx Bordeaux (project EcoGenPin).

## Data availability

All data will be made available in the Dryad repository (https://datadryad.org) as soon as the manuscript has been accepted.

## Supporting Information

**Table S1.1** Number of genotypes from the CLONAPIN clonal common garden used to study adaptive traits in *Pinus pinaster*.

**Figure S2.1** Phenological stages of bud burst.

**Protocol S3.1** Laboratory protocol for *Diplodia sapinea* inoculation.

**Figure S3.1** Pictures of the four scales of needle discoloration found along the necrosis caused by artificial *Diplodia sapinea* inoculation on excised branches of *Pinus pinaster*.

**Protocol S3.2** Laboratory protocol for *Armillaria ostoyae* inoculation.

**Table S4.1** MCMCglmm Bayesian model parametrization.

**Figure S5.1** Stripcharts of trait Best Linear Unbiased Predictors (BLUPs) for each genotype by population.

**Table S6.1 (see separate excel-file: TableS6.1.pdf)** All significant genotype effects (including additive, dominance and overdominance effects) of Single Nucleotide Polymorphisms (SNPs, minor allele frequency (MAF) > 0.1) on height, growth phenology and pathogen susceptibility traits identified by a two-step approach based on mixed-effects linear models (MLMs) implemented in Tassel and the Bayesian framework in BAMD (BMLMs). Bayesian mean SNP-effects and 95% credible intervals (CIs) were obtained from the distribution of the last 20,000 iterations in BAMD. Marker names and linkage groups (LG) as reported in Plomion *et al*. (2016). Site annotations: nc, non-coding (untranslated regions or introns); non-syn, non-synonymous; syn, synonymous; unk, unknown. *N*, number of phenotypic observations included in the analyses.

**Figure S6.1** Density plots of the effect sizes based on 20,000 BAMD simulations (left) and genotypic effects (box plots, right) for two non-synonymous Single Nucleotide Polymorphisms showing significant association with bud burst in 2017.

