## Supplementary_Table_S6.1 for "Genetic basis of growth, phenology and susceptibility to biotic stressors in maritime pine"

| | | | | | | | | Mixed linear model (TASSEL) | | | | | | | | | Bayesian framework (BAMD) | | | | | | | | | | | |
|---|---|---|---|---|---|---|---|---|---|---|---|---|---|---|---|---|---|---|---|---|---|---|---|---|---|---|---|---|
| | | | | | | | | | | | | | | | | | Additive model | | | Dominant model (allele 1) | | | Dominant model (allele 2) | | | Overdominance model | | |
| Trait | snp_id_inia | SNP name | SNP motif | site annotation | LG | MAF | N | F | p | additive effect | additive F | additive p | dominant effect | dominant F | dominant p | MarkerR$^2$ | Mean effect | (95% CIs) | | Mean effect | (95% CIs) | | Mean effect | (95% CIs) | | Mean effect | (95% CIs) | |
| *A. ostoyae* necrosis length | | AL750513_302 | [A/G] | nc | 1 | 0.3850 | 180 | 7.4765 | 0.0014 | -0.8298 | 8.0397 | 0.0066 | 0.9229 | 5.3819 | 0.0245 | 0.1041 | | | | 1.1731 | 0.4420 | 1.9091 | | | | 0.7189 | 0.0095 | 1.4034 |
| *A. ostoyae* necrosis length | | BX679585_950 | [A/G] | unk | 8 | 0.4397 | | 7.3967 | 0.0015 | 0.0830 | 0.0839 | 0.7732 | -1.5460 | 14.3500 | 0.0004 | 0.1007 | | | | -0.9544 | -1.7073 | -0.1991 | | | | -0.9965 | -1.7053 | -0.2912 |
| *A. ostoyae* necrosis length | | F51TW9001AI9YZ_1847 | [A/G] | unk | 7 | 0.2731 | | 5.9284 | 0.0048 | 1.3039 | 10.3043 | 0.0023 | -0.0881 | 0.0272 | 0.8697 | 0.0807 | -0.7349 | -1.3448 | -0.1294 | | | | -0.9402 | -1.8870 | -0.0152 | | | |
| *A. ostoyae* necrosis length | | F51TW9001ANBBN_100 | [T/C] | unk | 11 | 0.1679 | | 6.0467 | 0.0044 | -1.0199 | 6.5162 | 0.0137 | -0.4418 | 0.5753 | 0.4516 | 0.0823 | | | | | | | 0.8626 | 0.0088 | 1.7259 | | | |
| *A. ostoyae* necrosis length | | F51TW9001CXU1D_1264 | [T/C] | unk | 6 | 0.3644 | | 6.5943 | 0.0028 | 1.6422 | 12.7859 | 0.0008 | -0.8273 | 2.4751 | 0.1218 | 0.0898 | -0.9972 | -1.7872 | -0.2250 | -1.7246 | -3.0874 | -0.4280 | | | | | | |
| bb2015 | snp1167 | BX249218-322 | [A/C] | nc | | 0.315 | 411 | 6.49934 | 0.00167 | -1.07E+01 | 11.03273 | 9.77E-04 | -1.11E+01 | 6.39485 | 0.01183 | 0.02993 | 7.2298 | 2.10704 | 12.442 | | | | | | | 13.1179 | 3.44696 | 22.6922 |
| bb2015 | snp1304 | BX249671_307 | [T/C] | unk | 7 | 0.397 | | 6.43831 | 0.00177 | -1.05E+01 | 11.14411 | 9.21E-04 | 8.09829 | 4.11443 | 0.04317 | 0.02899 | 6.1292 | 0.76318 | 11.447 | 14.8503 | 5.61643 | 24.126 | | | | | | |
| bb2015 | snp1957 | BX252800-1728 | [A/C] | unk | 7 | 0.456 | | 6.02974 | 0.00263 | -3.85E+00 | 1.49247 | 0.22254 | -1.21E+01 | 9.30498 | 0.00243 | 0.02712 | | | | | | | -8.669 | -15.43646 | -1.9317 | 9.5705 | 1.74316 | 17.4099 |
| bb2015 | snp2124 | BX253890-151 | [T/G] | nc | 12 | 0.157 | | 6.35024 | 0.00192 | -1.71E+01 | 10.26034 | 0.00147 | -7.22E+00 | 1.19791 | 0.27438 | 0.02856 | 10.9449 | 4.29786 | 17.552 | 9.9087 | 1.98322 | 17.835 | | | | | | |
| bb2015 | snp2707 | BX681281_30 | [A/G] | unk | 1 | 0.254 | | 5.60635 | 0.00396 | -4.22E+00 | 1.34722 | 0.24645 | 16.119 | 10.88691 | 0.00105 | 0.02545 | | | | | | | 8.3426 | 1.0294 | 15.6906 | | | |
| bb2015 | snp2929 | CL2033CT1302CN1398-513 | [A/G] | nc | 1 | 0.408 | | 5.43621 | 0.00468 | -7.67E+00 | 6.33064 | 0.01225 | -5.66E+00 | 1.8744 | 0.17173 | 0.02451 | 7.1339 | 2.00113 | 12.321 | | | | -8.0104 | -14.84987 | -1.2437 | 10.342 | 3.05859 | 17.6062 |
| bb2015 | snp3100 | CL544Contig1_03.Pipn-84 | [T/G] | unk | | 0.135 | | 9.10696 | 1.35E-04 | -1.89E+01 | 10.74816 | 0.00113 | 2.69493 | 0.16005 | 0.68932 | 0.04097 | 10.1816 | 2.92232 | 17.579 | 16.801 | 0.61794 | 33.528 | -8.7781 | -16.99965 | -0.6334 | 12.1684 | 3.95323 | 20.4906 |
| bb2015 | snp3182 | CR392131-121 | [A/G] | unk | 3 | 0.491 | | 6.16228 | 0.00231 | -1.35E-01 | 0.00227 | 0.96206 | -1.34E+01 | 12.31459 | 5.00E-04 | 0.02779 | | | | | | | -9.3949 | -16.00276 | -2.8588 | 7.8448 | 0.09882 | 15.54 |
| bb2015 | snp4635 | F51TW9001BEJOH-703 | [G/C] | non-syn | 11 | 0.438 | | 5.52303 | 0.0043 | 1.87738 | 0.37815 | 0.53894 | -1.27E+01 | 9.82098 | 0.00185 | 0.0249 | | | | -10.1661 | -17.99729 | -2.286 | -8.8707 | -15.91143 | -1.8965 | | | |
| bb2015 | snp4798 | F51TW9001C6IZ8-79 | [T/A] | nc | 11 | 0.362 | | 5.62428 | 0.0039 | 7.34192 | 4.93349 | 0.02689 | 12.97593 | 9.2073 | 0.00257 | 0.02567 | | | | 12.9704 | 2.95593 | 23.069 | | | | | | |
| bb2015 | snp4914 | F51TW9001D5P2Y-1441 | [T/C] | non-syn | 5 | 0.214 | | 5.57156 | 0.0041 | 12.83801 | 8.48143 | 0.00379 | -1.63E+01 | 8.66034 | 0.00344 | 0.02543 | | | | -20.8939 | -34.86667 | -6.907 | | | | | | |
| bb2015 | snp5244 | FN692276-550 | [A/G] | unk | 12 | 0.402 | | 7.90304 | 4.29E-04 | -5.90E+00 | 3.87027 | 0.04983 | -1.18E+01 | 8.37638 | 0.00401 | 0.03555 | 5.6674 | 0.66379 | 10.774 | | | | -9.8216 | -16.81854 | -2.8406 | 10.4201 | 3.0625 | 17.8103 |
| bb2015 | snp5432 | i09773s1097 | [A/G] | non-syn | 12 | 0.194 | | 6.15419 | 0.00233 | -2.26E+00 | 0.28926 | 0.59099 | 17.10683 | 10.08025 | 0.00161 | 0.02768 | | | | | | | 11.9605 | 4.34136 | 19.6057 | -11.9063 | -19.52437 | -4.2665 |
| bb2015 | snp5605 | i13066s710 | [A/C] | nc | | 0.242 | | 6.10653 | 0.00244 | -1.42E+01 | 12.0129 | 5.84E-04 | -7.77E+00 | 2.15488 | 0.14289 | 0.02791 | 8.365 | 2.32783 | 14.407 | 8.3817 | 0.41851 | 16.334 | | | | 18.0323 | 5.63786 | 30.6181 |
| bb2015 | snp5677 | i16267s380 | [A/G] | unk | 2 | 0.411 | | 7.38793 | 7.05E-04 | 12.16869 | 14.51461 | 1.61E-04 | -2.07E+00 | 0.26406 | 0.60762 | 0.03331 | -10.5496 | -15.88515 | -5.168 | -12.5563 | -22.0943 | -2.945 | | | | -10.9217 | -18.31928 | -3.6878 |
| bb2015 | snp5717 | LP3-3-298 | [C/G] | unk | | 0.143 | | 5.53802 | 0.00424 | -2.23E+00 | 0.17982 | 0.67175 | 14.50341 | 4.97907 | 0.0262 | 0.02491 | 8.5102 | 1.69568 | 15.302 | 12.2695 | 4.08115 | 20.474 | 12.9459 | 4.74409 | 21.2559 | | | |
| bb2017 | snp0018 | 0_12730_01_contig1-159 | [A/C] | unk | 12 | 0.379 | 400 | 9.74934 | 7.36E-05 | -6.63E+00 | 17.03897 | 4.47E-05 | -1.49E+00 | 0.4026 | 0.52611 | 0.04388 | 4.00254 | 1.5091 | 6.4987 | 5.5809 | 1.36895 | 9.9685 | | | | 4.646579 | 0.797 | 8.5472 |
| bb2017 | snp0063 | 0_4105_01_contig2-279 | [A/G] | syn | 7 | 0.1 | | 7.78239 | 4.84E-04 | -1.07E+01 | 5.98503 | 0.01486 | -2.01E+01 | 14.79606 | 1.40E-04 | 0.03502 | | | | | | | -5.51405 | -10.7365 | -0.35039 | | | |
| bb2017 | snp0688 | AL750488_1167 | [A/C] | unk | | 0.189 | | 5.78574 | 0.00334 | 3.98526 | 2.41562 | 0.12093 | -5.13E+00 | 2.39476 | 0.12254 | 0.02611 | | | | -4.4435 | -8.57653 | -0.3351 | | | | | | |
| bb2017 | snp0743 | AL750545-695 | [T/A] | non-syn | 1 | 0.487 | | 5.7544 | 0.00344 | -4.93E+00 | 8.77821 | 0.00323 | 3.56754 | 2.43217 | 0.11967 | 0.0259 | 4.13065 | 1.37497 | 6.7996 | 5.1458 | 1.07883 | 9.1981 | | | | | | |
| bb2017 | snp0827 | AL750773_910 | [T/A] | unk | 3 | 0.499 | | 5.57962 | 0.00411 | 5.26758 | 8.90093 | 0.00305 | -3.13E+00 | 1.63618 | 0.20168 | 0.02618 | -4.04754 | -6.81041 | -1.339 | -5.0693 | -9.02944 | -1.125 | | | | | | |
| bb2017 | snp1098 | BX248967-202 | [G/C] | nc | 12 | 0.491 | | 6.45872 | 0.00174 | 5.93898 | 12.36493 | 4.88E-04 | -1.90E+00 | 0.704 | 0.40195 | 0.02908 | | | | -5.1898 | -9.35651 | -1.0206 | | | | | | |
| bb2017 | snp1337 | BX249816-2143 | [A/G] | non-syn | 7 | 0.269 | | 5.78336 | 0.00335 | -6.06E+00 | 6.48868 | 0.01124 | 1.00208 | 0.11285 | 0.7371 | 0.02616 | 5.78712 | 2.79164 | 8.8394 | 7.443 | 3.86209 | 11.0763 | 6.444604 | 2.6242 | 10.29718 | 7.23199 | 1.2103 | 13.3402 |
| bb2017 | snp1794 | BX251999-509 | [A/T] | unk | 9 | 0.296 | | 6.10471 | 0.00245 | -1.87E+00 | 0.85477 | 0.35577 | -8.93E+00 | 11.50773 | 7.63E-04 | 0.02752 | | | | | | | -5.530118 | -9.3896 | -1.60181 | | | |
| bb2017 | snp2548 | BX678760-1291 | [A/G] | unk | | 0.357 | | 5.82092 | 0.00322 | 4.61829 | 5.90167 | 0.01557 | 3.2465 | 1.68041 | 0.19562 | 0.0262 | | | | | | | 5.557376 | 0.4181 | 10.77013 | -5.162547 | -9.1268 | -1.1954 |
| bb2017 | snp2707 | BX681281_30 | [A/G] | unk | 1 | 0.254 | | 5.86913 | 0.00308 | -5.55E+00 | 6.95232 | 0.0087 | 9.16527 | 10.41352 | 0.00136 | 0.02655 | | | | 9.6608 | 3.97617 | 15.3402 | 5.470574 | 1.5715 | 9.41973 | | | |
| bb2017 | snp2969 | CL2640CT2248CN2410-1340 | [A/C] | unk | 6 | 0.477 | | 6.21882 | 0.00219 | 5.17299 | 9.5545 | 0.00214 | 3.8351 | 2.93205 | 0.08762 | 0.02799 | | | | | | | | | | -4.799382 | -9.2512 | -0.4238 |
| bb2017 | snp3549 | CT576106-142 | [C/G] | unk | 1 | 0.18 | | 7.1471 | 8.93E-04 | -1.05E+01 | 13.92479 | 2.18E-04 | 6.16293 | 3.41176 | 0.06548 | 0.03222 | 5.58994 | 2.16499 | 9.0293 | 9.9691 | 3.0869 | 16.9083 | | | | | | |
| bb2017 | snp3629 | CT577217-282 | [T/C] | syn | 8 | 0.268 | | 6.14051 | 0.00237 | 3.58743 | 2.95929 | 0.08617 | -9.55E+00 | 12.27979 | 5.11E-04 | 0.02771 | | | | -5.4787 | -10.92815 | -0.2104 | | | | | | |
| bb2017 | snp3644 | CT577489-1569 | [A/C] | unk | | 0.203 | | 5.8257 | 0.00321 | 4.89381 | 4.22021 | 0.0406 | -3.36E+00 | 1.09873 | 0.29518 | 0.02622 | -4.6057 | -7.5586 | -1.6017 | -4.4247 | -8.17673 | -0.6702 | | | | -6.707547 | -13.1702 | -0.3994 |
| bb2017 | snp3854 | CT580838-1007 | [T/G] | non-syn | | 0.178 | | 6.19074 | 0.00225 | 9.32138 | 10.80397 | 0.0011 | 10.98021 | 9.71119 | 0.00196 | 0.02786 | | | | | | | | | | -8.811789 | -16.138 | -1.6458 |
| bb2017 | snp3949 | CT582408-242 | [T/C] | nc | 1 | 0.199 | | 5.48155 | 0.00448 | 8.08976 | 10.02922 | 0.00166 | -8.44E+00 | 7.03398 | 0.00832 | 0.02467 | | | | -9.2085 | -16.15951 | -2.3467 | | | | | | |
| bb2017 | snp4415 | F51TW9001AOS8U-342 | [T/C] | unk | 4 | 0.23 | | 6.41299 | 0.00182 | -5.36E-01 | 0.05479 | 0.81505 | -8.80E+00 | 8.84747 | 0.00311 | 0.02886 | | | | | | | | | | 5.258851 | 1.0038 | 9.5012 |
| bb2017 | snp4433 | F51TW9001AQZUF-985 | [A/G] | unk | 4 | 0.231 | | 7.13029 | 9.07E-04 | 7.90594 | 12.78931 | 3.92E-04 | 8.72201 | 8.89216 | 0.00304 | 0.03209 | | | | | | | | | | -7.786754 | -14.1404 | -1.2995 |
| bb2017 | snp4526 | F51TW9001B1U5X-203 | [C/G] | nc | 12 | 0.301 | | 6.10794 | 0.00244 | 5.03265 | 6.34804 | 0.01215 | -8.54E+00 | 10.49017 | 0.0013 | 0.02758 | | | | -8.4096 | -13.80051 | -3.0379 | | | | | | |
| bb2017 | snp4622 | F51TW9001BD1TJ-1356 | [A/G] | non-syn | 7 | 0.301 | | 6.61478 | 0.00149 | 3.49988 | 3.06153 | 0.08094 | -9.16E+00 | 11.98503 | 5.95E-04 | 0.02977 | | | | -6.2492 | -11.23872 | -1.2385 | -6.644823 | -10.8296 | -2.37697 | | | |
| bb2017 | snp4837 | F51TW9001CGV5K-406 | [T/C] | syn | 1 | 0.49 | | 5.66907 | 0.00374 | -1.86E+00 | 1.20021 | 0.27395 | -7.45E+00 | 10.28621 | 0.00145 | 0.02556 | | | | | | | -5.451016 | -9.2519 | -1.66055 | 5.909973 | 1.6183 | 10.2801 |
| bb2017 | snp5184 | F7JJN6E01B7BCW-157 | [A/G] | syn | 5 | 0.117 | | 5.59136 | 0.00403 | -9.61E+00 | 4.32919 | 0.0381 | -1.55E+00 | 0.08885 | 0.7658 | 0.02516 | 7.70792 | 3.80051 | 11.5328 | 8.9058 | 4.79465 | 13.051 | 8.030913 | 3.5038 | 12.46737 | | | |
| bb2017 | snp5210 | FM945796-840 | [T/G] | unk | | 0.213 | | 6.28421 | 0.00206 | 8.82465 | 11.83887 | 6.42E-04 | -4.64E+00 | 2.06413 | 0.15159 | 0.02828 | -4.61252 | -7.74786 | -1.4726 | -9.0647 | -16.03004 | -2.2011 | | | | | | |
| bb2017 | snp5420 | i08906s326pg | [A/C] | unk | 8 | 0.195 | | 5.5134 | 0.00435 | -3.27E+00 | 1.34365 | 0.24709 | 10.54477 | 10.17113 | 0.00154 | 0.02481 | | | | | | | 6.166005 | 1.7452 | 10.4568 | | | |
| bb2017 | snp5496 | i10996s1211 | [T/C] | unk | | 0.301 | | 8.28116 | 3.00E-04 | -8.66E+00 | 16.54437 | 5.73E-05 | 5.43781 | 3.89869 | 0.04902 | 0.03727 | 4.29486 | 1.46655 | 7.1489 | 8.1132 | 2.37365 | 13.879 | | | | 4.353295 | 0.5509 | 8.1653 |
| bb2017 | snp5515 | i11276s420 | [T/C] | unk | | 0.3 | | 5.4351 | 0.0047 | -5.08E+00 | 6.16149 | 0.01347 | 2.23844 | 0.70437 | 0.40183 | 0.02453 | | | | 6.2341 | 2.332 | 10.1236 | | | | | | |
| bb2017 | snp5739 | PFK-39 | [T/C] | unk | 12 | 0.154 | | 8.82899 | 1.77E-04 | -1.18E+01 | 13.46922 | 2.76E-04 | 14.46243 | 15.41273 | 1.02E-04 | 0.03973 | | | | | | | 6.216364 | 1.7283 | 10.65858 | | | |
| bb2017 | snp6049 | sp_v3.0_unigene4029-1461 | [T/G] | nc | 1 | 0.458 | | 5.59726 | 0.00401 | 1.30223 | 0.68169 | 0.4095 | -7.11E+00 | 9.91932 | 0.00176 | 0.02519 | | | | -4.8037 | -8.69304 | -0.9636 | -6.657552 | -10.3571 | -2.95392 | | | |
| *D. sapine* a needle disc. | snp2558 | BX679001-1418 | [T/C] | non-syn | 7 | 0.192 | 144 | 5.54434 | 0.00481 | 0.09734 | 10.02625 | 0.00189 | -5.49E-02 | 2.10568 | 0.14897 | 0.04948 | -0.054124 | -0.10245 | -0.00648 | | | | | | | | | |
| *D. sapinea* needle disc. | snp3372 | CT574726_30 | [T/C] | unk | | 0.435 | | 5.53943 | 0.00483 | -5.74E-02 | 11.06425 | 0.00112 | -1.61E-02 | 0.48755 | 0.48617 | 0.04944 | -0.0511914 | -0.08943 | -0.0135 | | | | | | | | | |
| *D. sapinea* needle disc. | snp5499 | i11062s233 | [G/C] | syn | 7 | 0.293 | | 5.596 | 0.00458 | -5.73E-02 | 9.66144 | 0.00228 | 0.0195 | 0.67194 | 0.41376 | 0.04994 | -0.0480514 | -0.08951 | -0.00671 | | | | | | | | | |
| *D. sapinea* necrosis length | snp0518 | AL750104-316 | [T/G] | unk | 1 | 0.292 | 139 | 6.73102 | 0.00164 | 0.05561 | 0.01418 | 0.9054 | 2.24962 | 11.56631 | 8.87E-04 | 0.03833 | | | | | | | 1.372576 | 0.4886 | 2.2492 | | | |
| *D. sapinea* necrosis length | snp0707 | AL750492_595 | [T/C] | unk | 1 | 0.121 | | 5.88671 | 0.00353 | -2.17E+00 | 11.5049 | 9.09E-04 | -1.17E+00 | 1.95518 | 0.16431 | 0.03308 | 0.89698 | 0.08095 | 1.71291 | | | | | | | 2.19581 | 0.614806 | 3.8075 |
| *D. sapinea* necrosis length | snp3466 | CT575341-960 | [A/C] | syn | | 0.238 | | 6.09874 | 0.00291 | -2.44E+00 | 11.43736 | 9.40E-04 | 1.57287 | 2.97843 | 0.08665 | 0.03427 | | | | 2.19507 | 0.38391 | 4.1308 | | | | 0.93064 | 0.002719 | 1.8631 |
| *D. sapinea* necrosis length | snp3712 | CT578306-640 | [T/G] | unk | | 0.4 | | 5.88752 | 0.00353 | -1.28E+00 | 8.9073 | 0.00337 | -4.96E-01 | 0.64827 | 0.42214 | 0.03308 | | | | | | | -0.983372 | -1.8325 | -0.1303 | 1.01072 | 0.114725 | 1.9071 |
| *D. sapinea* necrosis length | snp4139 | F51TW9001A3IDU-1407 | [T/C] | nc | | 0.232 | | 8.97727 | 2.18E-04 | 1.80878 | 13.93912 | 2.77E-04 | 2.1234 | 10.51753 | 0.00149 | 0.05045 | | | | | | | 1.095398 | 0.2548 | 1.9509 | -1.58471 | -2.910448 | -0.2644 |
| *D. sapinea* necrosis length | snp4512 | F51TW9001AYUPD-852 | [T/C] | non-syn | 3 | 0.153 | | 6.07568 | 0.00297 | 0.36988 | 0.36822 | 0.54499 | -2.68E+00 | 8.94859 | 0.0033 | 0.03414 | | | | | | | | | | 1.44443 | 0.040624 | 2.889 |
| *D. sapinea* necrosis length | snp4565 | F51TW9001B4VCH-246 | [T/C] | nc | | 0.422 | | 5.99456 | 0.0032 | 0.83283 | 4.10835 | 0.04463 | 1.32546 | 6.19653 | 0.01401 | 0.03369 | | | | | | | | | | -1.1806 | -2.104186 | -0.2608 |
| *D. sapinea* necrosis length | snp4566 | F51TW9001B4VCH-358 | [A/G] | syn | | 0.424 | | 5.99456 | 0.0032 | -8.33E-01 | 4.10835 | 0.04463 | 1.32546 | 6.19653 | 0.01401 | 0.03369 | | | | 1.71629 | 0.73172 | 2.7048 | | | | | | |
| *D. sapinea* necrosis length | snp4976 | F51TW9001EIZX5-362 | [A/G] | non-syn | | 0.476 | | 7.13735 | 0.00113 | -1.75E-01 | 0.21109 | 0.64664 | 2.04979 | 14.11176 | 2.55E-04 | 0.04011 | | | | | | | 1.131242 | 0.3159 | 1.9395 | | | |
| *D. sapinea* necrosis length | snp5037 | F51TW9002FT2ZF-1060 | [T/G] | unk | 12 | 0.485 | | 8.61384 | 3.01E-04 | 1.79351 | 14.52857 | 2.09E-04 | -9.68E-01 | 2.89263 | 0.09129 | 0.04844 | -1.2788 | -1.9916 | -0.57355 | -1.65613 | -2.79484 | -0.5326 | | | | | | |
| *D. sapinea* necrosis length | snp5349 | FN695885-1909 | [G/C] | nc | 5 | 0.279 | | 6.09312 | 0.00292 | -8.61E-01 | 2.74274 | 0.1 | 2.39445 | 12.17118 | 6.54E-04 | 0.03424 | | | | | | | 1.290174 | 0.4293 | 2.163 | | | |
| *D. sapinea* necrosis length | snp5478 | i10796s1462pg | [A/G] | nc | 12 | 0.408 | | 7.30542 | 9.69E-04 | 0.11345 | 0.08622 | 0.76949 | 2.02659 | 14.54974 | 2.06E-04 | 0.04105 | | | | | | | 1.503466 | 0.7398 | 2.2719 | | | |
| *D. sapinea* necrosis length | snp6042 | sp_v3.0_unigene36670-2103 | [T/G] | unk | 1 | 0.139 | | 5.61162 | 0.00455 | 0.74419 | 1.70618 | 0.19369 | -1.99E+00 | 10.34751 | 0.00162 | 0.03153 | | | | | | | -1.08883 | -1.9513 | -0.2256 | 2.06097 | 0.867181 | 3.3016 |
| height | snp0130 | 2_9280_01.Pipn-311 | [T/C] | unk | 5 | 0.211 | 425 | 6.01494 | 0.00266 | -4.07E+00 | 4.0596 | 0.04456 | -3.05E+00 | 1.38355 | 0.24017 | 0.02172 | | | | | | | -4.50365 | -7.96414 | -1.0999 | 4.71698 | 1.0246 | 8.39501 |
| height | snp1269 | BX249583-420 | [A/G] | unk | 1 | 0.266 | | 7.38327 | 7.06E-04 | 7.02276 | 13.55641 | 2.62E-04 | 6.46156 | 8.1421 | 0.00454 | 0.02676 | 3.95552 | 1.16211 | 6.7615 | 6.51611 | 0.98159 | 12.00424 | | | | 3.88641 | 0.40449 | 7.37663 |
| height | snp1794 | BX251999-509 | [A/T] | unk | 9 | 0.296 | | 6.57564 | 0.00154 | 6.00013 | 12.5744 | 4.35E-04 | 5.35649 | 6.01951 | 0.01455 | 0.02375 | 2.74859 | 0.26157 | 5.2554 | 5.58878 | 0.42891 | 10.77043 | | | | | | |
| height | snp2012 | BX253184_476 | [T/C] | unk | 1 | 0.219 | | 7.15101 | 8.93E-04 | 7.30774 | 12.67163 | 4.18E-04 | 7.97994 | 9.09792 | 0.00273 | 0.02765 | | | | 6.90336 | 1.73922 | 12.0496 | | | | | | |
| height | snp2906 | CL1694Contig1_04-201 | [T/G] | nc | 9 | 0.169 | | 6.54276 | 0.00159 | 6.61274 | 8.02144 | 0.00484 | 10.52853 | 12.70727 | 4.06E-04 | 0.02363 | | | | 7.79369 | 0.84573 | 14.76311 | 3.88817 | 0.25382 | 7.567 | | | |
| height | snp3063 | CL3736CT1511CN1615-81 | [A/G] | nc | 5 | 0.45 | | 6.14879 | 0.00233 | 3.24786 | 5.13986 | 0.02389 | -5.69E+00 | 8.60636 | 0.00353 | 0.02221 | | | | | | | -4.07257 | -7.29985 | -0.8327 | | | |
| height | snp3190 | CR392177-1312 | [A/G] | nc | 7 | 0.214 | | 5.77307 | 0.00337 | -4.52E+00 | 4.71438 | 0.0305 | 9.02281 | 11.5427 | 7.48E-04 | 0.02152 | | | | 6.43025 | 0.33558 | 12.36425 | 3.3084 | 0.085 | 6.5664 | | | |
| height | snp3304 | CT2714-442 | [T/A] | unk | | 0.381 | | 5.68304 | 0.00367 | 4.79423 | 11.03879 | 9.70E-04 | -2.96E+00 | 2.31052 | 0.12925 | 0.02052 | -2.91978 | -5.33519 | -0.5026 | | | | | | | | | |

| | | | | | | | | | | | | | | | | | | | | | | | | | | | | |
|---|---|---|---|---|---|---|---|---|---|---|---|---|---|---|---|---|---|---|---|---|---|---|---|---|---|---|---|---|
| height | snp3563 | CT576335-234 | [G/C] | nc | 4.1 | 0.492 | | 6.20778 | 0.0022 | -2.48E+00 | 3.24456 | 0.07237 | 5.54546 | 9.15651 | 0.00263 | 0.02242 | | | | 4.5096 | 0.94068 | 8.04895 | 3.95202 | 0.87636 | 7.0324 | | | |
| height | snp3783 | CT579373-506 | [A/C] | unk | 6 | 0.344 | | 5.83467 | 0.00317 | -4.10E+00 | 7.24298 | 0.0074 | 6.05375 | 8.96035 | 0.00292 | 0.02107 | | | | 5.16082 | 0.66455 | 9.64825 | 3.16079 | 0.03538 | 6.2687 | | | |
| height | snp3819 | CT580064-331 | [A/G] | unk | | 0.442 | | 6.82994 | 0.0012 | 3.98293 | 8.68246 | 0.00339 | 5.37162 | 7.7132 | 0.00573 | 0.02467 | | | | 5.47739 | 1.53601 | 9.45258 | | | | | | |
| height | snp4010 | CT583593-733 | [T/C] | nc | | 0.231 | | 6.88456 | 0.00114 | 0.40269 | 0.04643 | 0.8295 | 7.37717 | 9.13977 | 0.00265 | 0.02486 | | | | | | | 4.61466 | 1.15071 | 8.0526 | -4.22167 | -7.7613 | -0.7033 |
| height | snp4514 | F51TW9001AZG2W-933 | [C/G] | unk | 4 | 0.438 | | 9.99082 | 5.77E-05 | -5.19E+00 | 12.77114 | 3.93E-04 | 5.52176 | 8.1563 | 0.0045 | 0.03608 | 3.48046 | 0.93066 | 6.0833 | 5.80766 | 1.9511 | 9.67196 | 3.82383 | 0.63739 | 7.0266 | | | |
| height | snp4579 | F51TW9001B6MAF-1134 | [T/G] | nc | 4 | 0.27 | | 5.52307 | 0.00429 | 0.92263 | 0.28759 | 0.59205 | 7.10426 | 9.81231 | 0.00185 | 0.01995 | | | | | | | 3.35429 | 0.12624 | 6.5937 | | | |
| height | snp4820 | F51TW9001CDDD5-305 | [A/G] | non-syn | | 0.407 | | 5.43567 | 0.00467 | 0.46016 | 0.09654 | 0.75617 | 6.15415 | 9.85425 | 0.00181 | 0.01964 | | | | | | | 3.90067 | 0.71773 | 7.0742 | | | |
| height | snp4926 | F51TW9001DBXSZ-1463 | [C/G] | nc | 5 | 0.182 | | 6.67453 | 0.0014 | 5.55085 | 5.60456 | 0.01836 | 8.28917 | 8.50414 | 0.00373 | 0.02411 | | | | 8.59014 | 2.73647 | 14.53301 | | | | | | |
| height | snp5938 | sp_v3.0_unigene17345-1191 | [T/G] | nc | 9 | 0.344 | | 5.97676 | 0.00276 | 5.58183 | 11.60649 | 7.20E-04 | 3.98453 | 3.68666 | 0.05552 | 0.02159 | 3.23163 | 0.5875 | 5.8714 | 6.19414 | 1.3924 | 11.0915 | | | | | | |
| height | snp5962 | sp_v3.0_unigene18547-188 | [T/A] | unk | 8 | 0.399 | | 6.00924 | 0.00267 | -3.83E+00 | 7.325 | 0.00708 | 5.60678 | 8.41437 | 0.00392 | 0.0217 | | | | 6.74767 | 2.54485 | 10.88275 | | | | | | |
| dbb2015 | snp1253 | BX249539-1987 | [A/G] | unk | | 0.125 | 411 | 7.44364 | 6.68E-04 | -2.64E+01 | 9.9355 | 0.00174 | -3.70E+01 | 14.85356 | 1.35E-04 | 0.03469 | | | | | | | | | | 38.1806 | 14.9228 | 62.4313 |
| dbb2015 | snp1254 | BX249539-2285 | [T/C] | unk | 3 | 0.116 | | 7.62441 | 5.61E-04 | 26.44826 | 10.00455 | 0.00168 | -3.81E+01 | 15.20195 | 1.13E-04 | 0.03553 | | | | -38.0685 | -67.49 | -10.4485 | | | | | | |
| dbb2015 | snp2130 | BX253931-1781 | [A/G] | unk | 12 | 0.295 | | 6.95165 | 0.00107 | -4.74E+00 | 1.30729 | 0.25356 | 19.88625 | 13.20333 | 3.15E-04 | 0.0324 | | | | 23.9899 | 9.459 | 38.6131 | | | | -15.5285 | -24.141 | -6.7834 |
| dbb2015 | snp2962 | CL2507CT3369CN3610-193 | [T/A] | nc | 1 | 0.387 | | 5.40629 | 0.00482 | -1.15E+01 | 10.20012 | 0.00151 | 7.35642 | 2.34563 | 0.12641 | 0.0252 | 8.9814 | 2.20314 | 15.73013 | 17.5328 | 5.783 | 29.1214 | | | | | | |
| dbb2015 | snp4228 | F51TW9001A8FFG-171 | [A/G] | unk | | 0.133 | | 5.39865 | 0.00485 | 14.36234 | 4.47049 | 0.0351 | -2.72E+01 | 10.67564 | 0.00118 | 0.02523 | | | | | | | | | | | | |
| dbb2015 | snp4738 | F51TW9001BWV4H-219 | [T/C] | non-syn | | 0.462 | | 5.94129 | 0.00286 | -1.16E+01 | 10.11137 | 0.00159 | -1.06E+01 | 4.58573 | 0.03283 | 0.02769 | 7.1009 | 0.45786 | 13.72349 | | | | | | | 17.8397 | 6.1079 | 29.425 |
| dbb2015 | snp4914 | F51TW9001D5P2Y-1441 | [T/C] | non-syn | 5 | 0.214 | | 5.88072 | 0.00304 | 11.52799 | 5.00428 | 0.02583 | -2.20E+01 | 11.53869 | 7.49E-04 | 0.02786 | | | | -20.5613 | -38.327 | -2.7225 | -12.437 | -22.65778 | -2.1787 | | | |
| dbb2015 | snp5036 | F51TW9002FPGRE-170 | [A/G] | nc | | 0.346 | | 5.50975 | 0.00435 | -1.11E+01 | 8.35726 | 0.00405 | -1.20E+01 | 5.59988 | 0.01843 | 0.02568 | 9.4671 | 2.50699 | 16.5795 | | | | | | | 19.8462 | 8.0295 | 31.6203 |
| dbb2017 | snp0405 | AL749850_679 | [A/G] | unk | | 0.402 | 425 | 5.44933 | 0.00461 | 6.40085 | 6.68616 | 0.01005 | -7.19E+00 | 5.08893 | 0.02459 | 0.02053 | -4.9924 | -9.31762 | -0.6539 | -8.1557 | -15.25068 | -1.09975 | | | | | | |
| dbb2017 | snp1743 | BX251734-1732 | [T/C] | syn | 5 | 0.395 | | 7.69335 | 5.23E-04 | 2.25799 | 0.92947 | 0.33555 | -1.11E+01 | 13.20704 | 3.13E-04 | 0.02902 | | | | -6.1761 | -12.09225 | -0.25111 | -10.44744 | -15.93544 | -5.05479 | | | |
| dbb2017 | snp2386 | BX667542-94 | [T/C] | nc | | 0.201 | | 5.70974 | 0.00357 | 8.86335 | 8.51651 | 0.00371 | 10.52142 | 6.94256 | 0.00873 | 0.02151 | | | | | | | 7.83105 | 1.33421 | 14.29244 | | | |
| dbb2017 | snp2733 | BX681656-960 | [A/C] | syn | 5 | 0.481 | | 8.04967 | 3.71E-04 | 8.39634 | 15.6222 | 9.07E-05 | -6.68E-01 | 0.05356 | 0.81709 | 0.03036 | -6.4158 | -10.15376 | -2.6686 | -8.7311 | -14.39133 | -3.04592 | | | | | | |
| dbb2017 | snp3396 | CT574915-594 | [T/C] | unk | 5 | 0.19 | | 5.66593 | 0.00373 | -1.04E+01 | 9.8247 | 0.00184 | 10.8614 | 6.68642 | 0.01005 | 0.02135 | | | | 15.8061 | 5.6085 | 26.12396 | 6.91587 | 0.31851 | 13.49681 | | | |
| dbb2017 | snp3819 | CT580064-331 | [A/G] | unk | | 0.442 | | 6.2361 | 0.00214 | 6.91226 | 11.14231 | 9.19E-04 | 5.32606 | 3.21918 | 0.0735 | 0.02355 | -4.363 | -7.93375 | -0.8224 | | | | | | | | | |
| dbb2017 | snp3957 | CT582680-451 | [A/C] | unk | | 0.201 | | 7.38323 | 7.05E-04 | 11.05093 | 11.91634 | 6.12E-04 | 0.06603 | 3.06E-04 | 0.98605 | 0.02782 | -10.4807 | -15.54122 | -5.4695 | -10.4499 | -20.15762 | -0.75108 | 6.37838 | 0.32975 | 12.39479 | -8.966898 | -15.63683 | -2.5032 |
| dbb2017 | snp4514 | F51TW9001AZG2W-933 | [C/G] | unk | 4 | 0.438 | | 7.77653 | 4.83E-04 | -4.39E+00 | 3.80293 | 0.05183 | 10.43542 | 12.37654 | 4.82E-04 | 0.02935 | | | | 8.1368 | 1.80043 | 14.51163 | 9.14651 | 3.82957 | 14.44137 | | | |
| dbb2017 | snp4606 | F51TW9001BAW7V-405 | [A/G] | unk | 12 | 0.163 | | 9.0883 | 1.37E-04 | -1.26E+01 | 13.74587 | 2.37E-04 | 2.54343 | 0.33561 | 0.56268 | 0.0343 | 7.2867 | 2.66474 | 11.8542 | 13.442 | 2.50983 | 24.17218 | | | | 8.948851 | 2.89749 | 14.9566 |
| dbb2017 | snp5287 | FN694219-1268 | [T/C] | nc | 3 | 0.278 | | 6.62828 | 0.00146 | -8.95E+00 | 10.86134 | 0.00106 | 0.66172 | 0.03731 | 0.84694 | 0.025 | | | | | | | | | | 7.428535 | 1.70196 | 13.143 |
| dbb2017 | snp5688 | i17647s350pg | [G/C] | unk | | 0.157 | | 5.84686 | 0.00313 | -7.83E+00 | 4.65378 | 0.03155 | -2.91E+00 | 0.44164 | 0.50669 | 0.02203 | 5.7467 | 0.87803 | 10.6106 | | | | | | | 8.061028 | 2.01287 | 14.0843 |
