## Supplementary_Material for "Genetic basis of growth, phenology and susceptibility to biotic stressors in maritime pine"

**S1 Samples included in this study from the CLONAPIN clonal common garden in Cestas, Bordeaux region (France)**

**Table S1.1** Number of genotypes from the CLONAPIN clonal common garden used to study adaptive traits in *Pinus pinaster*. bb, bud burst; dbb, duration of bud burst; proces, processionary moth nest counts; necrosis, necrosis length; disc, needle discoloration; n#, number; AF, Atlantic France; AI, Atlantic Iberia; C, Corsica; CS, Central Iberia; SI, Southeastern Iberia; M, Morocco.

| Gene pool | Population | Latitude | Longitude | Altitude (m) | height | bb 2015 | dbb 2015 | bb 2017 | dbb 2017 | proces | *D. sapinea* necrosis | *D. sapinea* disc | *A. ostoyae*. necrosis |
|---|---|---|---|---|---|---|---|---|---|---|---|---|---|
| | | | | | n# genotypes | | | | | | | | |
| AF | Pleucadec | 47.78 | -2.34 | 80 | 18 | 18 | 18 | 17 | 18 | 18 | | | |
| AF | St. Jean des Monts | 46.76 | -2.03 | 6 | 23 | 23 | 23 | 23 | 23 | 23 | 23 | 22 | 10 |
| AF | Olonne sur Mer | 46.57 | -1.83 | 13 | 19 | 19 | 19 | 19 | 19 | 19 | | | |
| AF | Le Verdon sur Mer | 45.55 | -1.09 | 11 | 21 | 21 | 21 | 18 | 21 | 21 | | | |
| AF | Hourtin | 45.18 | -1.15 | 26 | 21 | 21 | 21 | 21 | 21 | 21 | | | |
| AF | Mimizan | 44.13 | -1.30 | 37 | 16 | 16 | 16 | 14 | 16 | 16 | 16 | 15 | |
| AF | Pétrocq | 44.06 | -1.30 | 31 | 19 | 19 | 19 | 19 | 19 | 19 | | | |
| AI | Lamuño | 43.56 | -6.22 | 134 | 8 | 8 | 8 | 8 | 8 | 8 | | | |
| AI | Puerto de Vega | 43.55 | -6.63 | 121 | 7 | 7 | 7 | 5 | 7 | 7 | | | |
| AI | Cadavedo | 43.54 | -6.42 | 210 | 10 | 10 | 10 | 9 | 9 | 10 | 9 | 9 | |
| AI | Sierra de Barcia | 43.53 | -6.49 | 240 | 6 | 6 | 6 | 5 | 6 | 6 | | | |
| AI | Castropol | 43.50 | -6.98 | 391 | 10 | 10 | 10 | 10 | 10 | 10 | | | |
| AI | Armayán | 43.31 | -6.46 | 498 | 8 | 8 | 8 | 8 | 8 | 8 | | | |
| AI | Alto de la Llama | 43.28 | -6.49 | 503 | 7 | 7 | 7 | 6 | 7 | 7 | | | |

| | | | | | | | | | | | | | |
|---|---|---|---|---|---|---|---|---|---|---|---|---|---|
| AI | Sergude | 42.82 | -8.45 | 298 | 21 | 21 | 21 | 19 | 21 | 21 | | | |
| AI | San Cipriano de Ribaterme | 42.12 | -8.36 | 300 | 7 | 7 | 7 | 6 | 7 | 7 | | | |
| AI | Leiria | 39.78 | -8.96 | 20 | 19 | 19 | 19 | 17 | 19 | 19 | 19 | 18 | 10 |
| C | Pinia | 42.02 | 9.47 | 10 | 14 | 14 | 14 | 14 | 14 | 14 | 14 | 13 | 10 |
| C | Pineta | 41.97 | 9.04 | 750 | 7 | 7 | 7 | 7 | 7 | 7 | 7 | 7 | |
| CI | San Leonardo | 41.84 | -3.06 | 1096 | 10 | 10 | 10 | 10 | 10 | 10 | | | |
| CI | Bayubas de Abajo | 41.52 | -2.88 | 998 | 19 | 19 | 19 | 19 | 19 | 19 | | | |
| CI | Cuellar | 41.38 | -4.48 | 830 | 25 | 25 | 25 | 20 | 25 | 25 | | | |
| CI | Coca | 41.26 | -4.5 | 800 | 14 | 14 | 14 | 13 | 14 | 14 | 14 | 14 | 10 |
| CI | Carbonero el Mayor | 41.17 | -4.28 | 845 | 6 | 6 | 6 | 6 | 6 | 6 | | | |
| CI | Valdemaqueda | 40.52 | -4.31 | 890 | 8 | 8 | 8 | 8 | 8 | 8 | | | |
| CI | Cenicientos | 40.28 | -4.49 | 1100 | 5 | 5 | 5 | 5 | 5 | 5 | | | |
| SI | Olba | 40.17 | -0.62 | 1002 | 16 | 16 | 16 | 14 | 16 | 16 | 15 | 14 | |
| SI | Boniches | 39.99 | -1.66 | 1104 | 6 | 6 | 6 | 6 | 6 | 6 | | | |
| SI | Quatretonda | 38.97 | -0.36 | 435 | 15 | 15 | 15 | 13 | 15 | 15 | | | |
| SI | Oria | 37.53 | -2.35 | 1223 | 23 | 23 | 23 | 23 | 23 | 23 | 23 | 23 | 10 |
| SI | Cómpeta | 36.83 | -3.95 | 903 | 4 | 4 | 4 | 4 | 4 | 4 | | | |
| M | Tamrabta | 33.60 | -5.02 | 1758 | 9 | 9 | 9 | 9 | 9 | 9 | 9 | 9 | 8 |
| | Total number of genotypes | | | | 429 | 422 | 422 | 403 | 428 | 429 | 151 | 146 | 60 |
| | Total number of trees | | | | 3204 | 3146 | 3152 | 1392 | 1846 | 3204 | 453 | 438 | 180 |

**S2 Phenological stages of bud burst**

**Figure S2.1** Phenological stages of bud burst: 0) bud without elongation, as at the end of winter, 1) elongation of the bud, 2) emergence of brachyblasts, 3) brachyblast begin to space, 4) elongation of the needles, 5) total elongation of the needles.

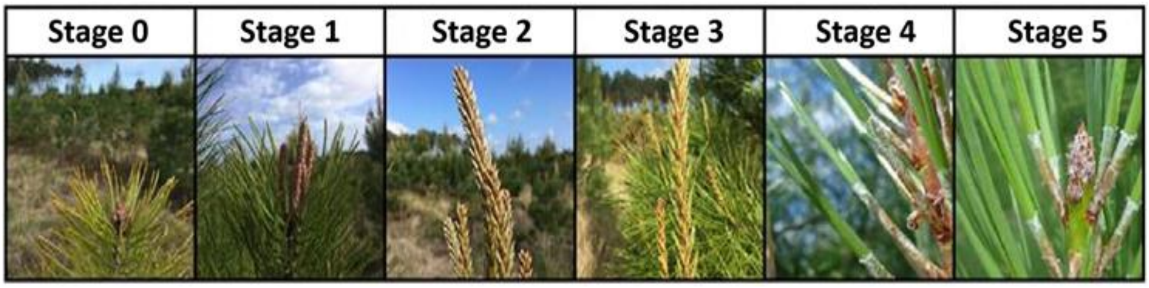

### S3 Pathogen inoculations on excised branches

#### S3.1 Laboratory protocol for *Diplodia sapinea* inoculation

1. Under the fume hood with sterilized material, *Diplodia sapinea* strain Pier4 was subcultured into 15 malt-agar Petri dishes. The pathogen was left to grow at room temperature for 3 days, during which it colonized the whole surface of the malt-agar.
2. The colonized Petri dishes were kept at 4°C to stop growth.
3. Shoots were collected in the CLONAPIN common garden, the phenological stage was estimated and the diameter was measured with a caliper.
4. We removed a needle fascicle in the middle of each shoot with a scalpel, making a small wound.
5. On the wound, we placed a 5mm diameter plug of malt-agar infected with *D. sapinea*, the mycelial side of the plug on the wound.
6. To keep the plug in place, we carefully wrapped the shoot in 3cm-wide cellophane.
7. The shoots were placed each in a glass jar with water, and kept in a climatic chamber set at 20°C with a daily cycle of 12h of light and 12 of dark.
8. Six days after inoculation, we removed the cellophane and the plug. The length of the necrosis around the wound was measured with a caliper, and needle discoloration was estimated from 0 – no discoloration to 3 – all needles discoloured along the necrosis. Other observations, such as "resin at the inoculation point" and "necrosis reaches the bud" were made but not used in the analysis of this study.

**Figure S3.1** Pictures of the four scales of needle discoloration found along the necrosis caused by artificial *Diplodia sapinea* inoculation on excised branches of *Pinus pinaster*. 0) No discoloration along the necrosis 1) Up to 50% of the needles are partly of fully discoloured 2) More than 50% of the needles are partly or fully discoloured 3) All needles are discoloured along the necrosis.

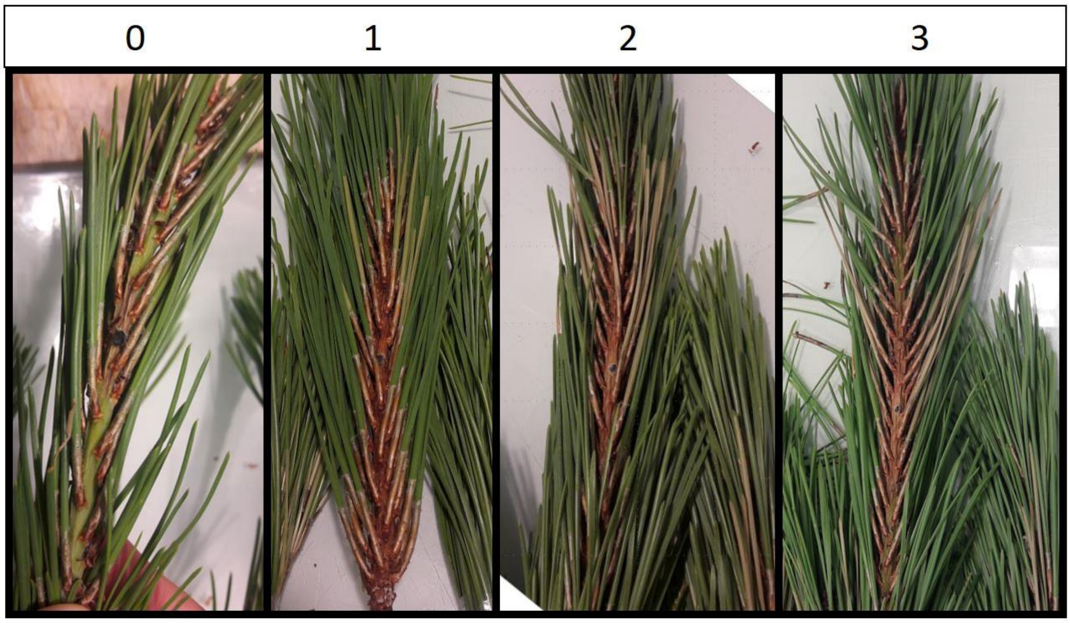

### S3.2 Laboratory protocol for *Armillaria ostoyae* inoculation

1. For *A. ostoyae* inoculation, we modified the protocols developed by Heinzelmann & Rigling (2016) and Ford *et al.* (2017). The liquid part of the medium consisted of 50% industrial vegetable soup (Knorr® 9 légumes) and 50% malt diluted in water (10 g of malt for 500 mL of water).
2. The mix was sterilized for 20 minutes at 120°C.
3. The solid part of the medium consisted of fresh hazelnut wood, sampled in Cestas (Nouvelle-Aquitaine, France) and chipped with an outdoor chipper. The chips were sieved then put in Sterilsop® bags (Hartmann®), sterilized a first time at 120°C for 20 min, placed in a heat chamber at 40°C to dry for 2h, then sterilized and dried a second time the same way. The bags were not opened during this process.
4. Bamboo sticks of approximatively 8 cm length were cleaned following the same process (see step 3).
5. All material was sterilized under UV light for 15 min under the fume hood before use.
6. Sterile laboratory jars (Dutscher®, 180 mL) were filled with hazelnut chips placing one bamboo stick in the middle as a place holder for the excised branch. The remaining space was filled with liquid medium. The jars were sealed with sterile cotton, aluminum foil and tape and re-sterilized 20 minutes at 120°C.
7. When the medium was cold, we inoculated each jar with two plugs of malt-agar of 5 mm of diameter infected with *A.ostoyae* and closed the jar with a lid.
8. The inoculated jars where then placed in the dark with firmly closed lids. After 2 months, we could not observe significant growth of the fungus, and some of the jars had to be discarded because of penicillium contamination inside the jar.
9. The remaining jars were placed in heat chambers at 23°C, 80% humidity with loosely closed lids to allow oxygenation (1/4 turn opened) (Lung-Escarmant, oral communication). After one month and a half, 180 jars showed satisfactory growth of *A.ostoyae*.
10. For each jar, the lid was opened, penicillium contamination estimated (on a scale from 0-no contamination to 5-very contaminated). Contaminated jars were safe to use as penicillium was only present on the surface of the mycelial culture. The bamboo stick was removed.
11. The lid was pierced with a Ø15 mm drill, replaced on the jar, and a branch was put in place of the bamboo stick. All of the jars and branches were treated this way, then replaced in the heat chamber with the same settings and an additional 12 h cycle of light/dark. The branches had a minimum length of 10 cm, and as the maximum length to fit inside the heat chamber was 30 cm, some branches were shortened. The length of the branch on sampling day depended on the growth of the year.
12. After 3 weeks, observations were made. Invasion success of *A. ostoyae* in the jar was visually estimated on a scale from 0 (0%) to 5 (more than 80%). Humidity was visually estimated in each jar (from 0 – dry to 3 – very humid). Finally, we measured the total length of the branch, the length of the branch before needles implantation, the length of the necrosis, and the length of the mycelium under the bark, with a caliper.

### S4 Estimation of genetic variance and heritability using MCMCglmm

**Table S4.1** MCMCglmm Bayesian model parametrization. Psrf stands for Gelman-Rubin criterion Potential Scale Reduction Factor, a mesure of model convergence. Good convergence of models is expected for psrf <1.02. bb, bud burst; dbb, duration of bud burst; necrosis, necrosis length; disc, needle discoloration; proces, processionary moth nest counts.

| | **Trait distribution** | **Link function** | **Co-variable** | **Prior fixed effects** | **Prior random effects** | **Prior residuals** | **Nb of iterations** | **Burn-in** | **Thinning** | **psrf** |
|---|---|---|---|---|---|---|---|---|---|---|
| height | Normal | identity | _ | default prior | V=1; n=0.002 | V=1; n=0.002 | 950,000 | 50,000 | 500 | 1.004 |
| bb2015 | Normal | identity | _ | default prior | V=1; n=0.002 | V=1; n=0.002 | 950,000 | 50,000 | 500 | 1.004 |
| bb2017 | Normal | identity | _ | default prior | V=1; n=0.002 | V=1; n=0.002 | 950,000 | 50,000 | 500 | 1.004 |
| dbb2015 | Normal | identity | _ | default prior | V=1; n=0.002 | V=1; n=0.002 | 950,000 | 50,000 | 500 | 1.003 |
| dbb2017 | Normal | identity | _ | default prior | V=1; n=0.002 | V=1; n=0.002 | 950,000 | 50,000 | 500 | 1.003 |
| *A. ostoyae* necrosis | Normal | identity | humidity in the jar* | default prior | V=1; n=0.002 | V=1; n=0.002 | 950,000 | 50,000 | 500 | 1.003 |
| *D. sapinea* necrosis | Normal | identity | _ | default prior | V=1; n=0.002 | V=1; n=0.002 | 950,000 | 50,000 | 500 | 1.003 |
| *D. sapinea* disc | Binomial | probit | _ | gelman prior | V=1; n=0.002; alpha.mu=0; alpha.v=1000 | Variance fixed at 1 | 950,000 | 50,000 | 500 | 1.003 |
| proces | Binomial | logit | height | default prior | V=1; n=0.002; alpha.mu=0 ; alpha.v=1000 | Variance fixed at 1 | 950,000 | 50,000 | 500 | 1.008 |

*Qualitative trait with 3 levels of humidity: dry, medium and very humid

### S5 Best Linear Unbiased Predictors (BLUPs) of phenotypic traits

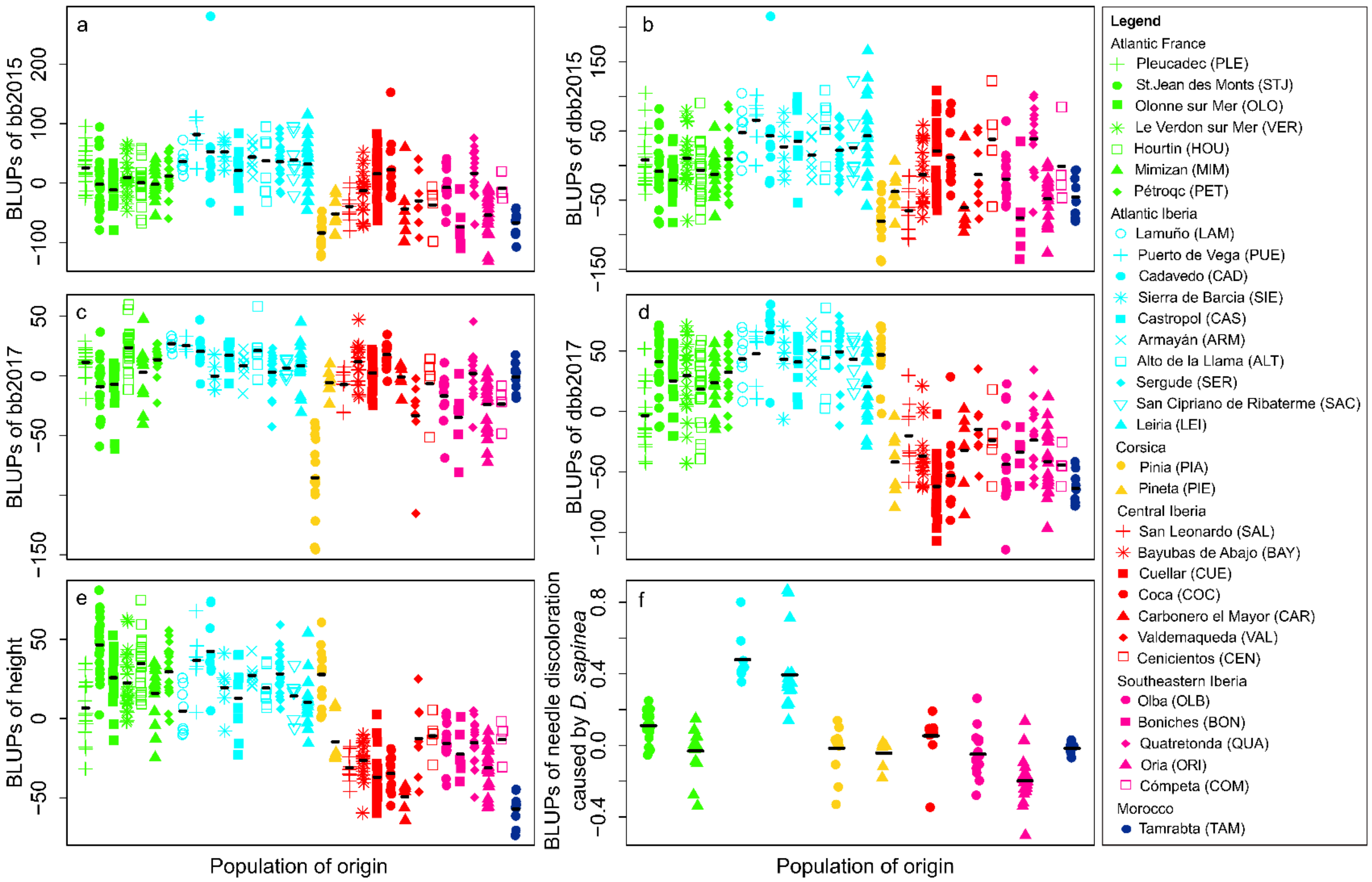


**Figure S5.1** Stripcharts of trait Best Linear Unbiased Predictors (BLUPs) for each genotype by population. Colours represent the gene pool (see Jaramillo-Correa, *et al.* 2015) and symbols represent the population in each gene pool (see legend). The black lines indicate the average BLUP value for each population. a) Bud burst (bb) in 2015, b) Duration of bud burst (dbb) in 2015, c) Bud burst in 2017, d) Duration of bud burst in 2017, e) Height, f) Needle discoloration caused by *Diplodia sapinea*. For necrosis length caused by *D. sapinea* and *A. ostoyae* see Figure 1 in main text.

### S6 Genotype-phenotype association

**Table S6.1 (see separate pdf-file: TableS6.1.pdf)** All significant genotype effects (including additive, dominance and overdominance effects) of Single Nucleotide Polymorphisms (SNPs, minor allele frequency (MAF) > 0.1) on height, growth phenology and pathogen susceptibility traits identified by a two-step approach based on mixed-effects linear models (MLMs) implemented in Tassel and the Bayesian framework in BAMD (BMLMs). Bayesian mean SNP-effects and 95% credible intervals (CIs) were obtained from the distribution of the last 20,000 iterations in BAMD. Marker names and linkage groups (LG) as reported in Plomion *et al*. (2016). Site annotations: nc, non-coding (untranslated regions or introns); non-syn, non-synonymous; syn, synonymous; unk, unknown. *N*, number of phenotypic observations included in the analyses.

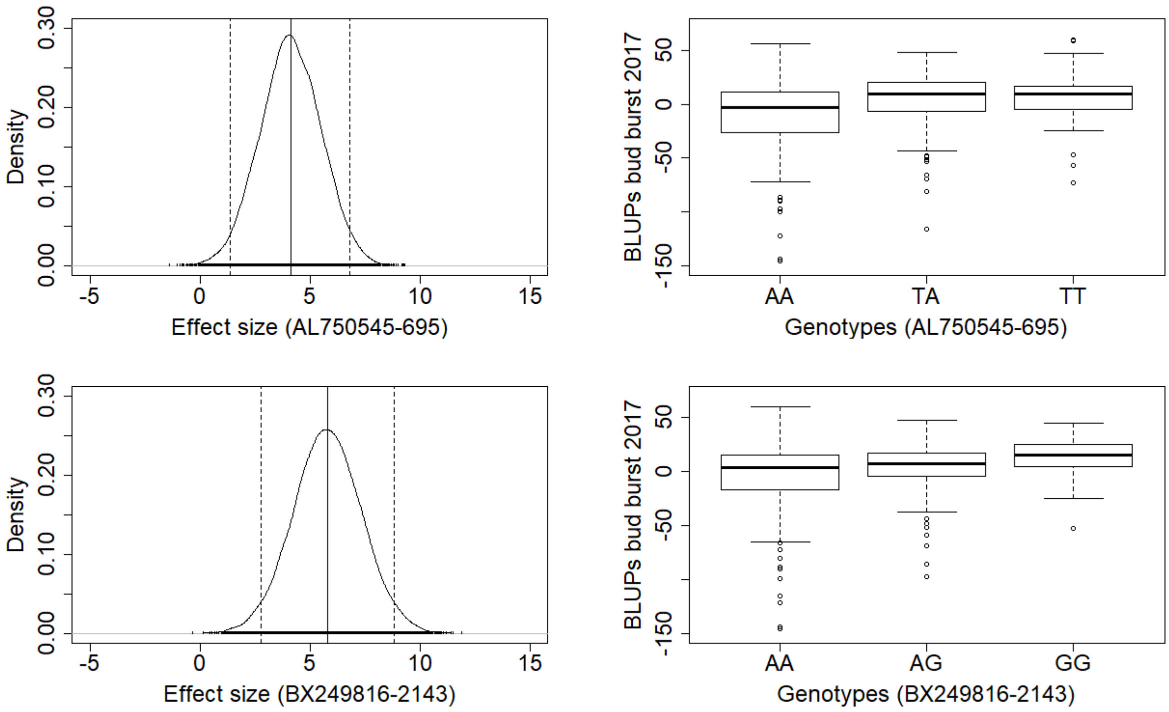


**Figure S6.1** Density plots of the effect sizes based on 20,000 BAMD simulations (left) and genotypic effects (box plots, right) for two non-synonymous Single Nucleotide Polymorphisms (SNPs, minor allele frequency (MAF) > 0.10) showing significant association with bud burst in 2017.
